# Targeted Degradation of Transcription Factors by TRAFTACs: Transcription Factor Targeting Chimeras

**DOI:** 10.1101/2020.10.12.336529

**Authors:** Kusal T. G. Samarasinghe, Saul Jaime-Figueroa, Katherine Dai, Zhenyi Hu, Craig M. Crews

## Abstract

Many diseases, including cancer, stem from aberrant activation and overexpression of oncoproteins that are associated with multiple signaling pathways. Although proteins with catalytic activity are able to be successfully drugged, the majority of other protein families, such as transcription factors, remain intractable due to their lack of ligandable sites. In this study, we report the development of TRAnscription Factor TArgeting Chimeras (TRAFTACs) as a generalizable strategy for targeted transcription-factor degradation. Herein, we show that TRAFTACs, which consist of a chimeric oligonucleotide that simultaneously binds to the transcription-factor of interest (TOI) and to HaloTag fused dCas9 protein, can induce degradation of the former via the proteasomal pathway. Application of TRAFTACs to two oncogenic TOIs, NF-κB and brachyury, suggests that TRAFTACs can be successfully employed for the targeted degradation of other DNA-binding proteins with minor changes to the chimeric oligonucleotide.

## Introduction

PROteolysis TArgeting Chimeras (PROTACs) are heterobifunctional molecules that target disease-causing proteins for proteasomal degradation (1, 2). PROTAC molecules simultaneously bind to both the target protein and a E3 ligase to induce proximity-dependent ubiquitination of the former, thereby tagging the target protein for proteasome-mediated degradation (2, 3). This technology was developed nearly two decades ago and has successfully been applied to cytoplasmic, nuclear proteins, and membrane-associated proteins (BCR-Abl and the receptor tyrosine kinases EGFR, and FLT3) (4–8). Targeted protein degradation offers several advantages over conventional small molecule-based inhibition including the potential to expand the druggable proteome to encompass non-catalytic, scaffolding proteins in deregulated biological systems. However, degradation of such proteins by PROTACs still requires a target ligand, and the development of ligands for proteins lacking a well-defined active site is challenging and time-consuming (9, 10).

TFs are DNA-binding proteins that directly or indirectly regulate gene expression and upon deregulation, are among the known drivers of pathologies. Therefore, many efforts have been devoted to therapeutically target TFs implicated in human diseases (11). TFs often mediate their regulatory functions through DNA and protein-protein interactions. Unlike kinases or other druggable proteins, development of small molecule inhibitors for TFs is truly challenging due to the lack of enzymatic activity and ligandable pockets. Therefore, development of alternative strategies to target such proteins is currently lacking. A small number of TFs, such as the estrogen and androgen receptors, which drive breast and prostate cancer, respectively, have been drugged by small molecules, although the emerging resistance to such therapies hinders their long-term effectiveness (12–15). Recently, these ligands were incorporated into PROTACs to induce the degradation of estrogen and androgen receptors: two PROTACs (ARV-471 and ARV-110) targeting estrogen and androgen receptors have already progressed to phase 1 clinical trials and demonstrated high tolerability and safety in humans (16–18). In addition, a selective STAT3 PROTAC was recently developed (19). However, steroid receptors and STAT3 are among the minority of TFs possessing a ligand binding site that controls their activity. Because many TFs mediate their regulatory functions through interactions with DNA and/or with other proteins, they frequently lack enzymatic activity and ligandable pockets—features that have been successfully exploited in developing small molecule inhibitors for more readily druggable proteins. Therefore, development of alternative, generalizable strategies to target TFs is paramount.

Since TFs mediate their transcriptional activity by interacting with DNA and other accessory proteins, a variety of protein-DNA interaction inhibitors and protein-protein interaction inhibitors (PPIs) have been developed (20). For instance, pyrrole-imidazole polyamide binds to the minor grove of DNA and inhibits binding of the pro-inflammatory transcription factor, NF-κB (21). In addition to DNA-protein interaction inhibitors, PPIs have also been extensively studied in TF inhibition. The transcription factor Myc forms a heterodimer with Max to bind to enhancer box sequences to regulate gene expression (22). Several groups have successfully developed PPIs that inhibit Myc/Max interaction, eliciting effects on gene expression (23–25). As a complementary strategy to these PPIs, stabilizing molecules of Max homodimers have also been developed to reduce the availability of Max to form transcriptionally active Myc/Max heterodimers (26, 27). The aforementioned approaches were directed at the TF themselves, but indirect approaches that block TF activity have also been reported. For example, specific STAT3/5 inhibitors have been developed to block nuclear STAT3 translocation by inhibiting its phosphorylation, dimerization and DNA binding (28, 29). The targeting of upstream effectors, another indirect approach to inhibit TFs, has also been well-studied (30, 31): for example, multiple NF-κB pathway inhibitors do so by targeting upstream signaling components, such as SRC/Syk, PI3k/Akt and IKK (32–34). BET protein targeting PROTACs that induced BRD4 degradation has also been shown to reduced downstream oncogenic c-Myc levels, leading to improved survival in mice (35, 36). These previous approaches to overcome the overexpression and/or hyperactivity of oncogenic TFs have motivated us to develop a strategy that bypasses the need for target protein ligand development while readily and directly targeting a target transcription-factor of interest (TOI).

In the current study, we have developed a generalizable strategy to induce TF degradation by co-opting the cellular degradation machinery. While development of traditional PROTACs for TF degradation is challenging due to the lack of cognate small molecule ligands, TFs are proteins that elicit their function via binding to specific DNA sequences. Therefore, by taking the advantage of this intrinsic DNA-binding ability of TFs, we developed chimeric oligos with TF-specific DNA sequences attached to an E3 ligase recruiting protein, and we termed these chimeric molecules, <u>TRA</u>nscription <u>F</u>actor <u>TA</u>rgeting <u>C</u>himera<u>s</u>, or **TRAFTACs**. Our chimeric oligo consists of a TOI-binding double stranded DNA covalently linked to a Cas9 CRISPR-binding RNA. The dsDNA binds to TOI, while the CRISPR-RNA binds to an ectopically-expressed dCas9HT7 (HT7-HaloTag7) fusion protein. An applied haloPROTAC then recruits the VHL-E3 ligase to the vicinity of the DNA-bound TOI via the complexed fusion protein, inducing ubiquitination and proteasomal degradation of TOI (Fig. 1). In this proof-of-concept study, we demonstrate that TRAFTACs can recruit both TOI and the E3-ligase via the intermediate protein dCas9HT7 and induce degradation of the disease-relevant TFs NF-κB and brachyury in the presence of haloPROTAC.

**Figure 1.**
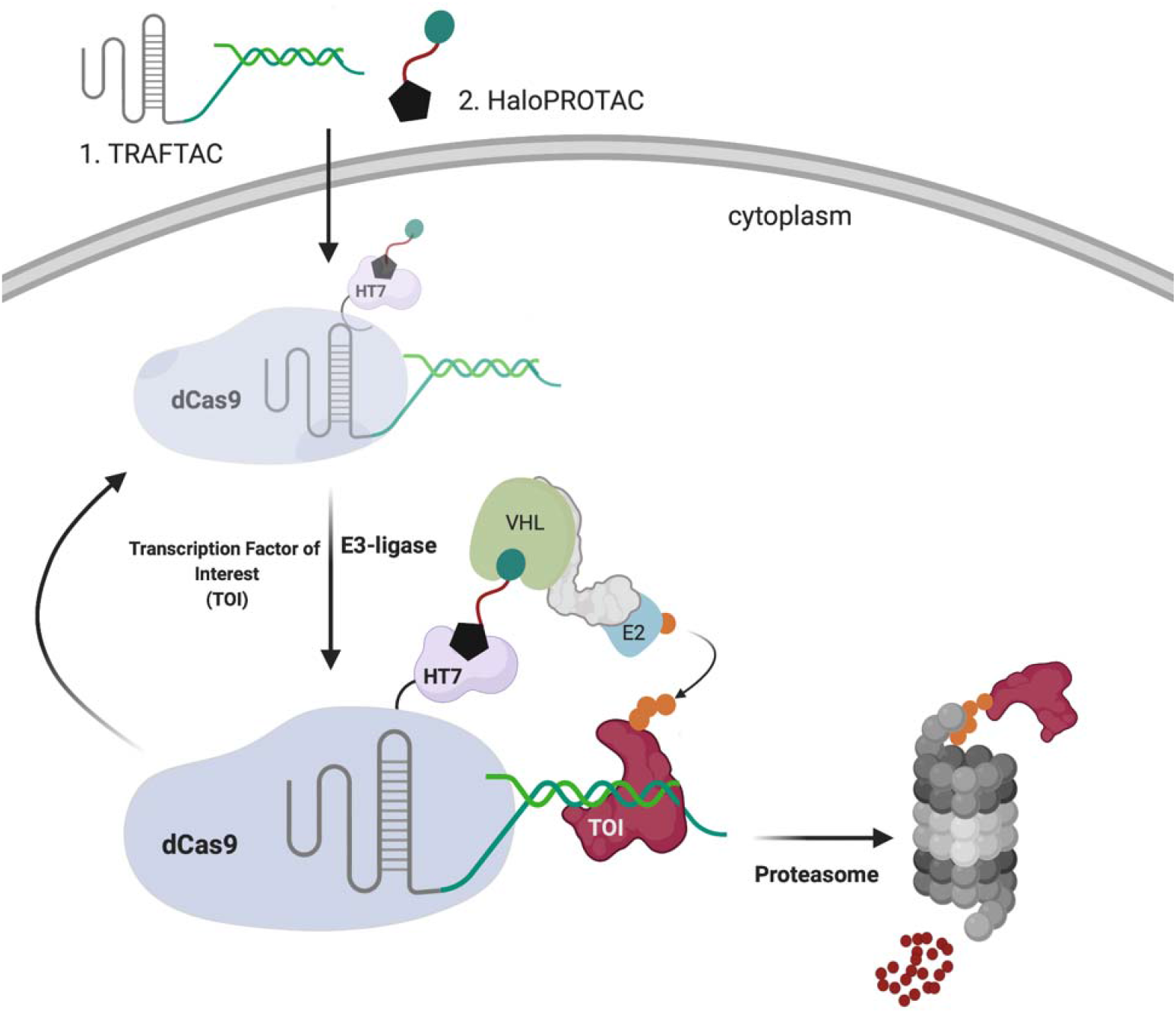
Schematic representation of the TRAnscription Factor TArgeting Chimeras (TRAFTACs). Heterobifunctional dsDNA/CRISPR-RNA chimera (TRAFTAC) recruits E3 ligase complex through dCas9-HT7 in the presence of haloPROTAC. Heterobifunctional TRAFTAC binds to dCas9-HT7 via its RNA moiety while dsDNA portion of the chimera binds to the transcription factor of interest (TOI). Addition of haloPROTAC recruits VHL E3 ligase complex to the vicinity of TOI. TRAFTAC-mediated proximity induced ubiquitination directs TOI for proteasomal degradation.

## Results

### NF-κB Engages with the dCas9HT7 Ribonucleocomplex

NF-κB binds to kappa B sequences upstream of its target genes (37). Therefore, we selected the kappa B sequence as the NF-κB binding element in our initial TRAFTAC design (Fig. 2A). A chimeric oligonucleotide containing the kappa B sequence at the 3’ end of a CRISPR RNA (crRNA) was synthesized. The double-stranded chimeric oligo (NFκB-TRAFTAC) was generated by annealing with the reverse complementary oligo of the kappa B sequence (Fig. S1A). Using an electrophoretic mobility shift assay (EMSA), we first evaluated the ability of crRNA to engage the dCas9HT7 fusion protein when the former is presented as a part of a double stranded chimeric oligo. The data indicated that NFκB-TRAFTAC engaged with purified dCas9HT7 fusion protein as the band is shifted upwards in the agarose gel (Fig. 2C and S2). Also, a single major band for NFκB-TRAFTAC indicated the stability of the chimeric molecule after the annealing reaction. Subsequently, we evaluated the interaction of the chimeric oligo with the fusion protein within cells. We transfected a fluorescein-labeled TRAFTAC into HEK293 cells that stably overexpress HA-tagged dCas9 fused at its C-terminus to HT7 (CT-dCas9HT7) (Fig. 2B). First, to test whether the NFκB-TRAFTAC successfully enters the cell, we analyzed cells by confocal microscopy after the transfection of fluorescein-labeled TRAFTAC (Fig. S1C). The intracellular green signal indicated that transfection of the chimeric NFκB-TRAFTAC was successful (Fig. S3). Next, we analyzed the cells by confocal microscopy to test whether NFκB-TRAFTAC can engage with dCas9HT7 in cells. After fixation, the cells were incubated with anti-HA primary antibody followed by Alexa fluor-568 labeled secondary antibody. The co-localization of fluorescein signal with immunofluorescence signal (Fig. 2D) suggested that dCas9HT7 fusion protein engages with NFκB-TRAFTAC in cells. Next, we tested for NF-κB (p50) engagement with the dCas9HT7 ribonucleocomplex (dCas9HT7:NFκB-TRAFTAC) by performing an immunoprecipitation experiment. Cell lysates were incubated with either NFκB-TRAFTAC or a control-TRAFTAC and then subjected to immunoprecipitation by HA agarose beads. The eluted samples were probed for p50 and HT7, and the results indicated that p50 binds to dCas9HT7 only in the presence of the NFκB-TRAFTAC (Fig. 2E). Overall, the data indicate that NFκB-TRAFTAC engages both the dCas9HT7 fusion protein and p50.

**Figure 2.**
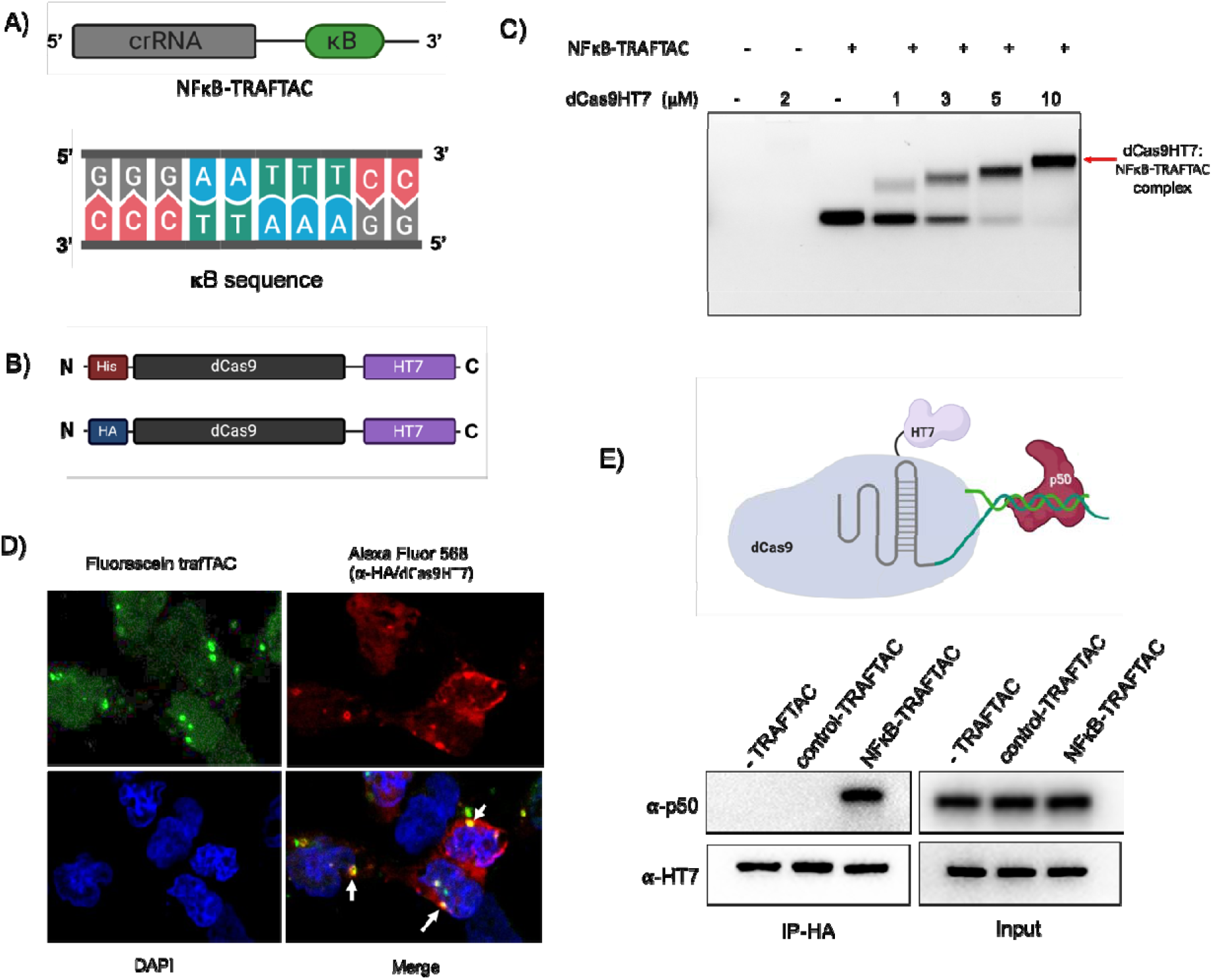
Binding experiments for NFκB-TRAFTAC, CT-dCas9HT7 and p50. A) A diagram illustrating the architecture of NF-κB-TRAFTAC and the NF-κB binding kappaB sequence. B) Schematic of bacterial and mammalian expression CT-dCas9HT7 constructs. After bacterial expression, protein was purified by His-tag affinity column purification. While the bacterial expression vector has a His-tag sequence, it was replaced by HA-tag for mammalian expression. C) Different concentrations of purified CT-dCas9HT7 were incubated with NFκB-TRAFTAC and the protein:oligo complexes separated on an agarose gel. Data represent the ability of dCas9HT7 to bind to the modified-double stranded chimeric NFκB-TRAFTAC. D) Immunofluorescence data illustrating *in cellulo* TRAFTAC engagement with fusion dCas9HT7. Cells were transfected with fluorescein labeled TRAFTAC for 12 h and cells fixed, permeabilized and probed with anti-HA antibody, followed by secondary antibody conjugated to Alexa Fluor 568. Images were captured using a confocal microscope. E) Immunoprecipitation of p50 and ribonucleocomplex (dCas9HT7:NFκB-TRAFTAC). Stable cells lysates with dCas9HT7 were treated with NFκB-TRAFTAC and control-TRAFTAC After 1 h, dCas9HT7 was immunoprecipitated using HA antibody conjugated beads and eluted samples were probed with antibodies as shown.

### TRAFTACs Induce NF-κB Degradation

In TRAFTAC design, we adapted a RNA-binding but catalytically dead CRISPR Cas9 (dCas9) protein as an intermediate spacer protein that brings the TOI and E3 complex into close proximity (38). Specifically, in our HEK293 cell line, we stably overexpressed CT-dCas9HT7, to simultaneously bind both the von-Hippel Lindau E3 ubiquitin ligase (VHL), via a haloPROTAC, and the TOI, through a TRAFTAC (Fig. 1). In previous studies, VHL-based haloPROTACs were designed to induce degradation of HaloTag fusion proteins (39). However, in this study, our intent is to employ HaloTag7 as a ligand inducible E3-recruiting element to cause TOI degradation but not dCas9HT7. Therefore, we first carried out an initial haloPROTAC screening using existing haloPROTACs and several newly synthesized haloPROTACs to select the optimal haloPROTAC that do not induce CT-dCas9HT7degradation. Our group’s previous study suggested that haloPROTACs with 3-PEG linkers are potent degraders of directly bound HaloTag7 fusion proteins, but that ability of haloPROTACs to induce degradation of a directly bound substrate protein was decreased with increasing linker lengths (39). Therefore, we synthesized several haloPROTACs with longer linker lengths and screened for their ability to induce CT-dCas9HT7 degradation. As anticipated, we observed that longer haloPROTACs are poor degraders of CT-dCas9HT7, providing a good starting point to test TRAFTACs (Fig. S4A). More specifically, haloPROTACs HP3, HP8 and HP10 with shorter linker lengths (3 and 4 PEG units) induced CT-dCas9HT7 degradation even at 2.5 μM, whereas HP13 (7 PEG units) and HP14 (9 PEG units) did not induce significant CT-dCas9HT7 degradation (Fig. S4A and S5). Two other, longer haloPROTACs, HP15 and HP16 (12 PEG units) were also tested for their ability to degrade CT-dCas9HT7 and consistently, we did not observe significant CT-dCas9HT7 degradation (Fig. S4B and S5). Therefore, we selected HP13, HP14, HP15 and HP16 as our haloPROTAC panel to test in TRAFTAC studies.

Before testing the ability of TRAFTACs to induce p50 degradation, we determined the minimum NFκB-TRAFTAC concentration that does not induce cellular cytotoxicity. The data obtained from MTS assays suggested that NFκB-TRAFTAC can be introduced to cells up to 50 nM without inducing significant cytotoxicity by lipofection (Fig. S6). To ensure lack of toxicity, we transfected a lower concentration (25 nM) of NFκB-TRAFTAC into CT-dCas9HT7 stable cells followed by the treatment of HP14 in the presence or absence of 5 ng/mL of TNF-alpha for 12 h. HP14 induced p50 degradation only in the presence of TNF-alpha (Fig. 3A). Latent NF-κB stays in the cytoplasm as an inactive heterotrimeric complex until extracellular stress signal is introduced. The inhibitor of Kappa B (IκB) binds to the NF-κB, physically masking its kappa B-binding sequence. However, TNF-alpha induces the release of active NF-κB from the inhibitory complex and facilitates TRAFTAC binding and subsequent degradation by NFκB-TRAFTAC. TNF-alpha treatment also induces the proteolytic processing of the NFκB precursor, p105 into p50 (40). Therefore, overall TRAFTAC accessible p50 levels are higher in the presence of TNF-alpha, compared to untreated cells (Fig. 3A, right panel). We also tested our panel of haloPROTACs to determine the effect of linker length in TRAFTAC-mediated p50 degradation. Interestingly while HP13 treatment induced p50 degradation to a similar extent as in HP14 treated cells, HP15 with 12 PEG unit linker did not induce p50 degradation (Fig. S8A). Left-handed haloPROTAC (HP16) with 12 PEG unit linker also did not induce p50 degradation.

**Figure 3.**
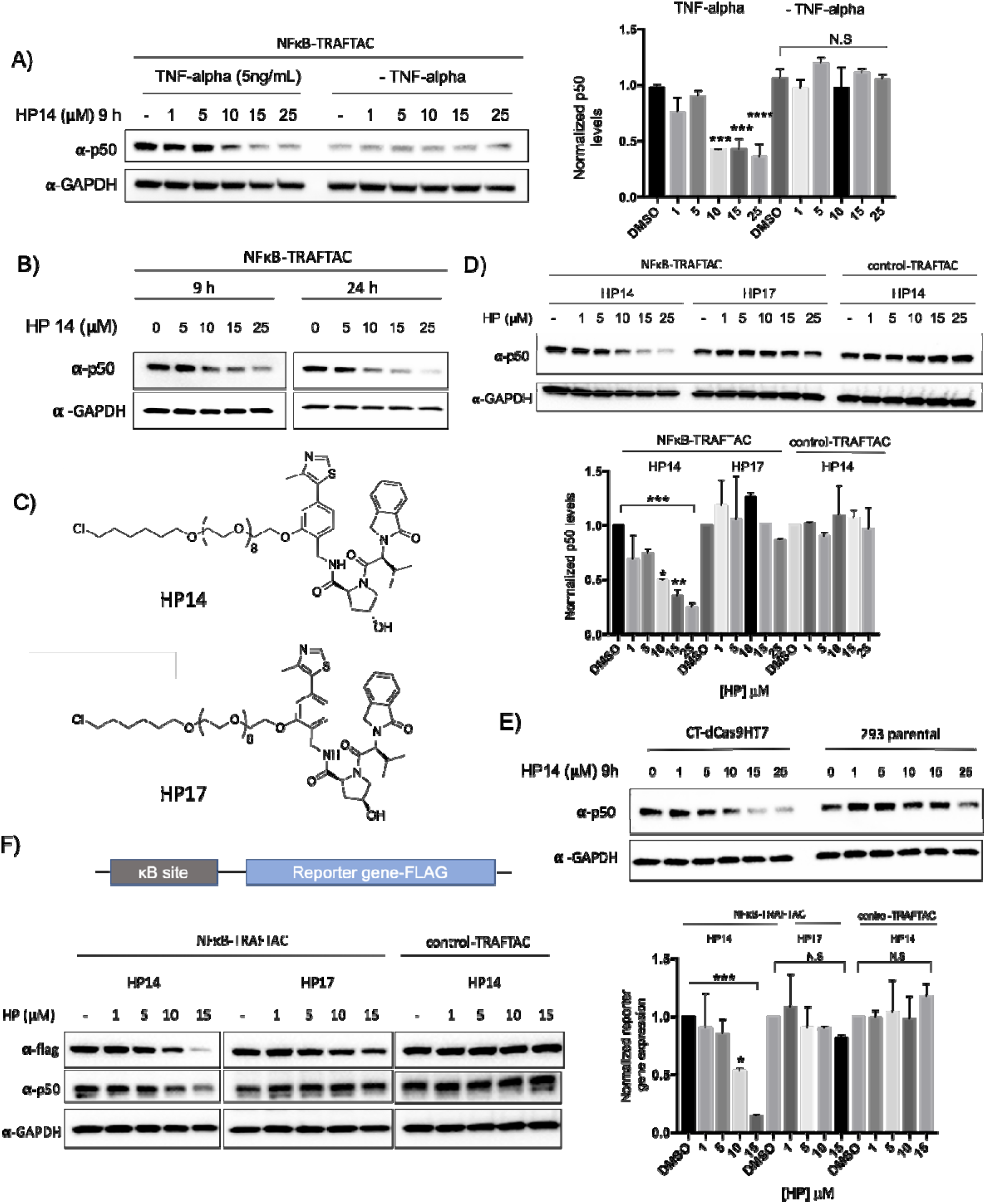
NF-κB degradation by TRAFTACs. A) HP14 induces p50 degradation in the presence of TNF alpha. Briefly, cells were transfected with NFκB-TRAFTAC and after 16 h, cells were treated with increasing HP14 concentrations. After 1h, cells were treated with or without TNF-alpha for indicated times. B) TRAFTACs induce p50 degradation within 9 h of HP14/TNF-alpha treatment. C) Chemical structures of HP14/TNF-alpha and the inactive epimer control (HP17). D) VHL and TRAFTAC-dependent p50 degradation. Cells were transfected with either with NFκB-TRAFTAC or control-TRAFTAC followed by the treatment of HP14/TNF-alpha or the HP17/ TNF-alpha for 12 h. Cell lysates were probed for p50 and GAPDH. E) TRAFTAC induced p50 degradation is dependent on dCas9HT7. Stable cells overexpressing dCas9HT7 and parental cells were transfected with chimeric oligo and the experiment performed as mentioned above. F) NFκB-TRAFTAC mediated p50 degradation elicit an effect on downstream gene expression. (Not significant (N.S); * P < 0.03; ** p<0.002; *** p<0.0002; **** p<0.0001; n=2)

Next, we evaluated TRAFTAC-mediated p50 degradation at both 9 and 24 h post HP14/TNF-alpha treatment. The data indicated that TRAFTACs could induce significant p50 degradation after 9 h of TNF-alpha treatment. However, the degradation levels of p50 only slightly improved over the 24 h of treatment (Fig. 3B). Moreover, TRAFTAC-mediated degradation was further intensified in the presence of protein synthesis inhibitor cycloheximide (CHX) demonstrating that TRAFTACs significantly lowered the half-life of p50 (Fig. S8B). To confirm whether the p50 degradation is dependent on VHL recruitment, we next compared TRAFTAC-induced degradation in response to HP14 to its inactive epimer control (HP17) (Fig. 3C). Gratifyingly, p50 degradation was observed only in the cells treated with HP14, whereas no significant p50 degradation was observed in HP17 treated cells (Fig. 3D left and center panel). Next, cells stably overexpressing CT-dCas9HT7 were transfected with NFκB-TRAFTAC or control-TRAFTAC (Fig. S1B). Cells were subsequently treated with HP14 for 12 h and cell lysates probed for p50 levels. As shown in Figure 3D, HP14 induced p50 degradation in NFκB-TRAFTAC transfected cells. In contrast, cells transfected with control-TRAFTAC did not induce significant loss of p50 levels, suggesting that TRAFTAC-mediated p50 degradation is dependent on the dsDNA portion of the TRAFTAC (Fig. 3D-right panel). In addition to p50 degradation, NFκB-TRAFTAC also induced the degradation of second NFκB heterodimeric subunit, RelA (p65) (Fig. S7B). Furthermore, parental cells, without CT-dCas9HT7 expression, did not induce p50 degradation, confirming that the observed target degradation is dependent on dCas9HT7 as well (Fig. 3E). Finally, we performed a reporter gene assay to test the effect of TRAFTAC-mediated p50 degradation on downstream gene expression. Cells stably expressing CT-dCas9HT7 were transiently transfected with a NF-κB reporter plasmid that expresses FLAG tagged reporter gene under the control of NF-kB. After TRAFTAC transfection and HP14 treatment, cell lysates were analyzed for reporter gene expression. The results demonstrated a loss in κB promoter activation that was consistent with the p50 degradation observed only in response to NFkB-TRAFTAC. Importantly, the control-TRAFTAC did not reduce reporter levels, nor did co-treatment with the inactive HaloPROTAC HP17. We observed that only HP14 co-treatment repressed reporter gene expression (Fig. 3F).

### HaloTag7 Position Relative to dCas9 Governs Its Susceptibility to Degradation

We had employed 2.5 μM of haloPROTACs in the initial screening to select the best haloPROTAC for TRAFTAC investigations. While HP14 did not induce p50 degradation at that concentration, our data indicated that significant p50 degradation occurred at concentrations over 10 μM HP14. However, HP14 also induced CT-dCas9HT7 degradation within that concentration range (Fig. 4A). Thus, we tested whether the observed p50 degradation was an indirect consequence of overall CT-dCas9HT7:NFκB-TRAFTAC:p50 complex degradation (Fig. S9A). This was accomplished using HP3, a short, potent, HT7 fusion protein HaloPROTAC (39), together with NFκB-TRAFTAC to test whether this combination also induces p50 degradation. We hypothesized that the shorter haloPROTAC (HP3) should be much more restrictive in terms of the steric range of protein ubiquitination induced by the recruited VHL. To this end, cells stably overexpressing CT-dCas9HT7 were transfected with NFκB-TRAFTAC followed by the treatment of either HP3 or HP14. Although HP3 could induce significant degradation of CT-dCas9HT7, it did not induce p50 degradation. In contrast, HP14 induced the degradation of both (Fig. 4B and S9B). Clearly, the data showed that ligand-mediated degradation of CT-dCas9HT7 does not mandate concomitant p50 degradation. Thus, HP14-mediated p50 degradation is not a result of CT-dCas9HT7 fusion protein degradation. Having successfully ‘’uncoupled” degradation of the two complexed proteins, we next set about engineering the reciprocal scenario: facilitation of p50 degradation while sparing degradation of dCas9HT7.

**Figure 4.**
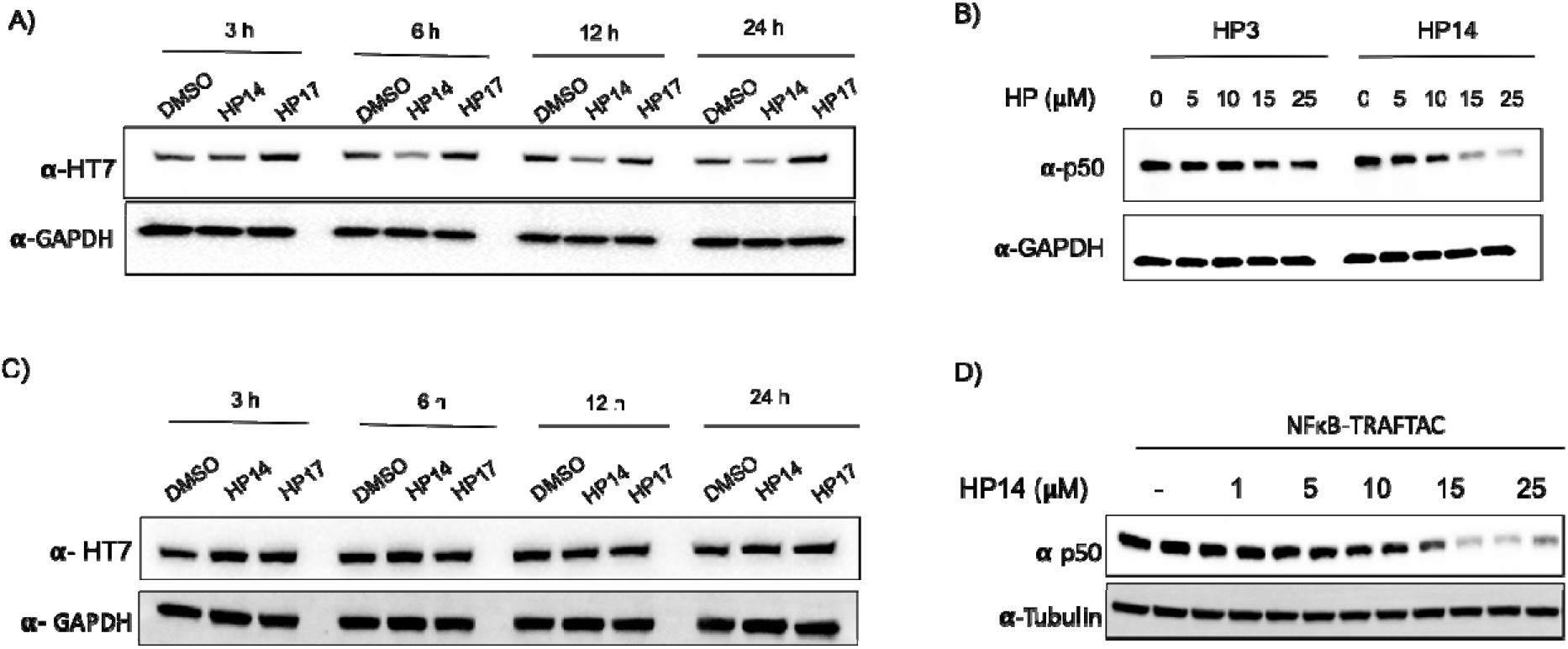
Positioning of HT7 in dCas9 governs the HT7 fusion protein degradation by haloPROTACs. A) HP14 induces degradation of CT-dCas9HT7 fusion protein. The cell line stably overexpressing CT-dCas9HT7 was treated with HP14 and HP17 and lysed at indicated time points. Lysates were probed for HT7 and GAPDH. B) The haloPROTAC HP3 (possessing a shorter linker) did not induce p50 degradation, in contrast to HP14. C) Cells stably expressing NT-dCas9HT7 were treated with HP14 and lysed at indicated time points. D) NFκB-TRAFTAC/HP14/TNF-alpha treated NT-dCas9HT cells were lysed and probed for p50 and tubulin levels.

To improve the TRAFTAC system accordingly, we generated a construct with HT7 at the N-terminus of dCas9 (NT-dCas9HT7). The positioning of susceptible lysine residues on a targeted protein of interest and the presentation and orientation of the E3 complex are important factors for a successful PROTAC-mediated degradation (39, 41–43). Therefore, we speculated that fusion of HaloTag7 at the opposite end of the dCas9 might alter the HP14-induced dCas9HT7 degradation. To investigate this, we generated cells stably overexpressing NT-dCas9HT7 protein and incubated them with increasing concentrations of HP14, followed by probing for NT-dCas9HT7 levels. Interestingly, we observed significantly less degradation levels of NT-dCas9HT7 by HP14 compared to that of CT-dCas9HT7 (Fig. 4C). Next we evaluated the same cell line to test whether NT-dCas9HT7 can be used to induce p50 degradation together with NFκB-TRAFTAC. First, we transfected NFκB-TRAFTAC followed by HP14 and TNF-alpha treatment for 9 h. Then cell lysates were probed for p50. The data showed that NT-dCas9HT7 can, indeed, induce p50 degradation (Fig. 4D). From the Cas9 crystal structure bound to guide RNA, it is apparent that the 3’ end of the guide RNA, the N-terminus and C-terminus of Cas9 are all positioned at the same face of the protein (PDB ID:4OO8; Fig. S10) (44). This remarkable positioning of the dCas9 and crRNA offers an explanation as to how the E3 ligase recruited via both NT- and CT-HaloTag7 can access NFκB-TRAFTAC-bound proteins and induce their degradation. Overall, the data suggest that dCas9 protein is less susceptible to HP14-mediated degradation when HaloTag7 is fused at the N-terminal end and therefore provides a better tool for use in TRAFTAC design. Therefore, we employed the NT-dCas9HT7 fusion protein (which retains the N-terminal HA tag) in subsequent experiments.

### TRAFTACs are Generalizable: TRAFTACs Induce Brachyury Degradation

Current TRAFTAC technology is based on the intrinsic ability of TFs to bind to specific DNA sequences. Therefore, we hypothesized that TRAFTACs should be adaptable to target a variety of transcription factors for proteasomal degradation by simply customizing the DNA sequence specific to a TOI. To investigate the generalizability of TRAFTACs to target different TFs, we extended our approach to induce the degradation of the T-box transcription factor T, also known as brachyury. Brachyury is a key transcription factor that regulates notochord formation in embryonic stages of development, and its downregulation leads to tail development defects (45, 46). Recent studies have also suggested that brachyury knockdown leads to the inhibition of tumorigenesis, cell migration and invasion, suggesting a key role in cancer metastasis (47). Furthermore, genomic amplification of the locus that harbors brachyury gene, mutations and germline tandem duplication of the brachyury gene are associated with cancer such as chordoma (48–51). Given the importance of brachyury in cancer, we sought to target brachyury for proteasomal degradation using the TRAFTAC technology. To this end, we synthesized a new chimeric oligo that comprised a dsDNA portion that recognizes and binds brachyury and the previously-used crRNA sequence capable of recruiting NT-dCas9HT7, the E3 ligase-recruiting intermediary-fusion-protein (Fig. 5A and S11C). First, we confirmed the binary (dCas9HT7: brachyury-TRAFTAC) and ternary (dCas9HT7: brachyury-TRAFTAC: brachyury) complex formation by EMSA experiments. Consistent with NFκB-TRAFTAC EMSA data, we showed that brachyury-TRAFTAC engages the dCas9HT7 fusion protein (Fig. S11A). Further, data suggest a successful ternary complex formation of dCas9HT7:brachyury-TRAFTAC:brachyury whereas a control-TRAFTAC failed to do so (Fig. 5C). Next, we performed an immunoprecipitation experiment to test whether brachyury could engage with the NT-dCas9HT7 ribonucleocomplex. We generated a cell line that stably overexpresses both brachyury-GFP and NT-dCas9HT7 proteins. The cells lysates were subjected to immunoprecipitation with HA antibody attached to agarose beads in the absence of a chimeric oligo or in the presence of control-TRAFTAC or brachyury-TRAFTAC. The eluates were probed for dCas9HT7 protein and brachyury-GFP using antibodies against HaloTag7 and brachyury respectively. The results indicated that brachyury selectively engages NT-dCas9HT7 protein only when brachyury-TRAFTAC is present, suggesting successful brachyury recruitment by the chimeric oligo to the dCas9HT7 fusion protein (Fig. 5B).

**Figure 5.**
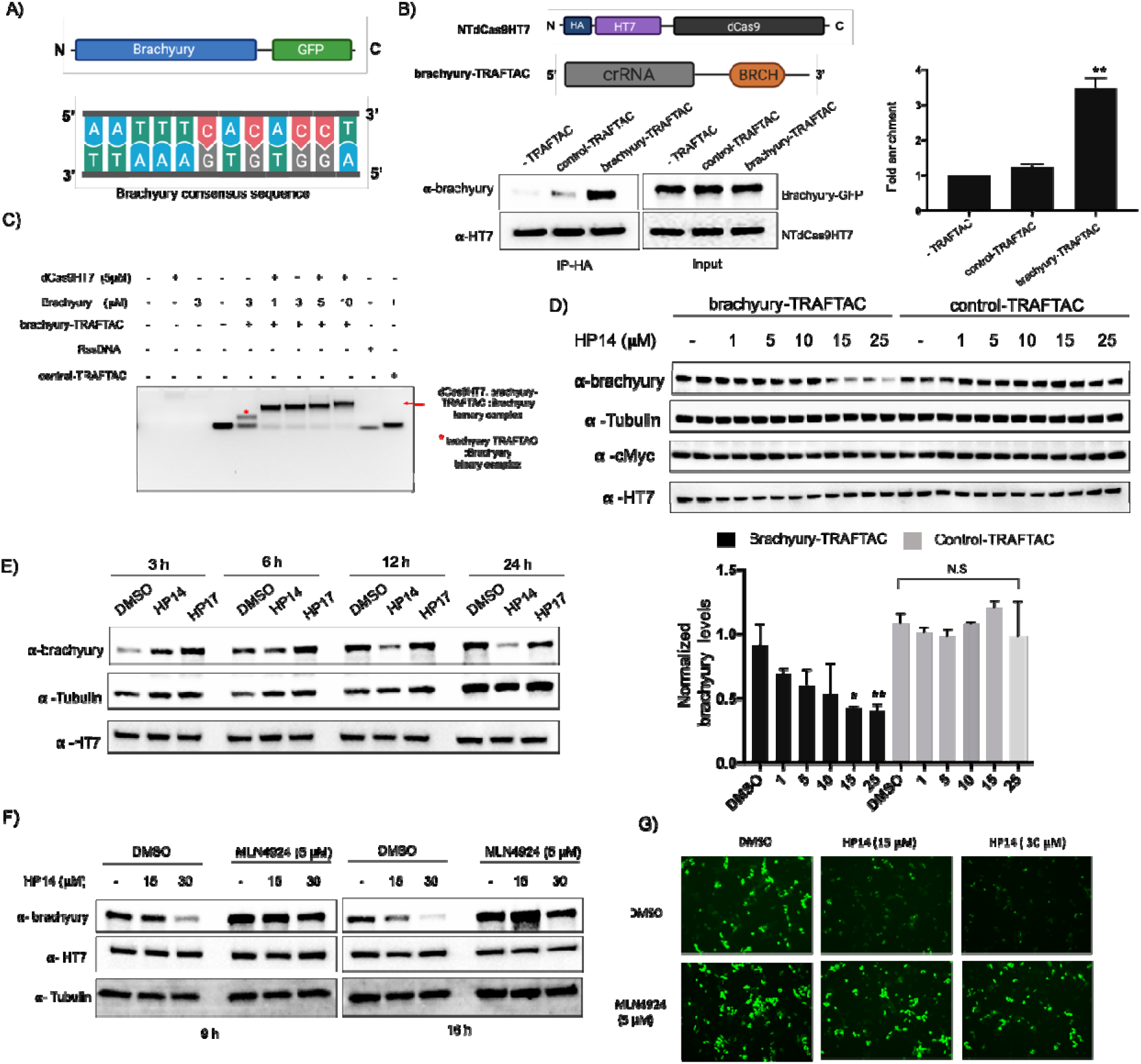
TRAFTACs targeting brachyury induces its degradation. A) Schematic representation of the brachyury-GFP construct and the brachyury binding DNA sequence. B) Cell lysates that overexpress brachyury-GFP were incubated with the brachyury-TRAFTAC and the inactive control-TRAFTAC before HA immunoprecipitation. Eluted samples were probed against brachyury and HT7 levels. C) Ternary complex formation assay for the dCas9HT7:chimeric oligo and brachyury. Purified proteins were incubated with chimeric oligos and protein:oligo complexes were separated using DNA agarose gel to analyze gel shifts as depicted in the figure. D) Cells stably expressing brachyury-GFP were transfected with brachyury-TRAFTAC or control-TRAFTAC followed by HP14 treatment. Cells were then lysed and analyzed for brachyury, tubulin, c-Myc and HT7 levels. (Not significant (N.S); * P < 0.03; ** p<0.002) E) Time dependent degradation of brachyury-GFP. Cells stably expressing brachyury-GFP were transfected with brachyury-TRAFTAC and treated with HP14 and HP17. Cells were lysed at indicated time points, subjected to western blotting and probed for brachyury, tubulin and HT7. F) Stable cells overexpressing brachyury-GFP were treated with HP14 and MLN4924 for 9 and 16 h. Cells were lysed and probed for brachyury, HT7 and tubulin. G) The Brachyury-GFP fluorescence signal was captured after HP14 and MLN4924 co-treatment.

To test whether brachyury-targeting TRAFTACs induce target degradation, cells stably expressing brachyury-GFP were transfected with brachyury-TRAFTAC or control-TRAFTAC, followed by the treatment with increasing HP14 concentrations. Western blot analysis indicated that brachyury-TRAFTAC induces brachyury-GFP degradation at low micromolar HP14 concentrations. Further, HP14 induced brachyury-GFP degradation within 12 h and reached maximum degradation after 24 h (Fig. 5E). The inactive epimer, HP17 did not induce brachyury-GFP degradation indicating that the observed degradation is dependent on VHL-E3 ligase recruitment (Fig. S11D). Similarly, the control-TRAFTAC did not induce brachyury-GFP degradation in response to HP14 suggesting that the observed degradation is also dependent on brachyury-TRAFTAC double stranded DNA (Fig. 5D). Additionally, HP14 co-treatment with a neddylation inhibitor, MLN4924, failed to induce brachyury-GFP degradation, further confirming that TRAFTAC induces brachyury degradation via the proteasomal pathway (Fig. 5F). We also investigated brachyury-GFP levels using fluorescence microscopy and observed GFP signal reduction, further confirming the observed degradation pattern in the western blot analysis (Fig. 5G). Overall, the data show that the brachyury-targeting TRAFTAC can successfully induce brachyury-GFP fusion protein degradation via the proteasome.

Next, we investigated the potential of TRAFTACs to induce proteasomal degradation of endogenous brachyury protein. Brachyury is predominantly expressed in early embryonic development. However, it has been shown that brachyury is also expressed in several cancer cell lines, including Hela cells. Therefore, we selected Hela cells to test TRAFTAC-mediated endogenous brachyury degradation. We first evaluated whether brachyury engages dCas9HT7 through brachyury-TRAFTAC, by performing an immunoprecipitation experiment using a cell line that expresses NT-dCas9HT7. The data indicate that brachyury successfully binds dCas9HT7 (Fig. 6A). The data obtained after the transfection of brachyury-TRAFTAC and treatment with HP14 at different concentrations indicated that TRAFTACs can also successfully degrade endogenous brachyury protein at concentrations of HP14 < 10 μM (Fig. 6B, left panel). We have confirmed that the observed endogenous brachyury degradation is VHL-dependent as the epimer control failed to induce brachyury degradation (Fig. 6B, right panel). Furthermore, the data obtained with the control-TRAFTAC oligo confirmed that TRAFTAC-induced endogenous-brachyury degradation is DNA-sequence dependent (Fig. S11B). To further confirm sequence dependency, we also tested the levels of other transcription factors. We analyzed p65 and c-Myc levels using the same experimental protocol and the data indicated that brachyury-targeting TRAFTAC did not affect either p65 or c-Myc levels (Fig. 6B and S12). This suggests that TRAFTAC-mediated degradation is selective and displays a minimal effect on other DNA binding proteins. Furthermore, we tested an all-scrambled version of the chimeric oligo, allscrambled-TRAFTAC, composed of both scrambled crRNA and scrambled dsDNA sequences (Fig. S1D). The western blot analysis showed that upon transfection of allscrambled-TRAFTAC, HP14 failed to induce brachyury degradation compared to brachyury-TRAFTAC transfected cells (Fig. 6C). Collectively, the data suggest that TRAFTAC-mediated degradation of endogenous brachyury is dependent on the specific dsDNA sequence, dCas9, VHL and the proteasomal pathway.

**Figure 6.**
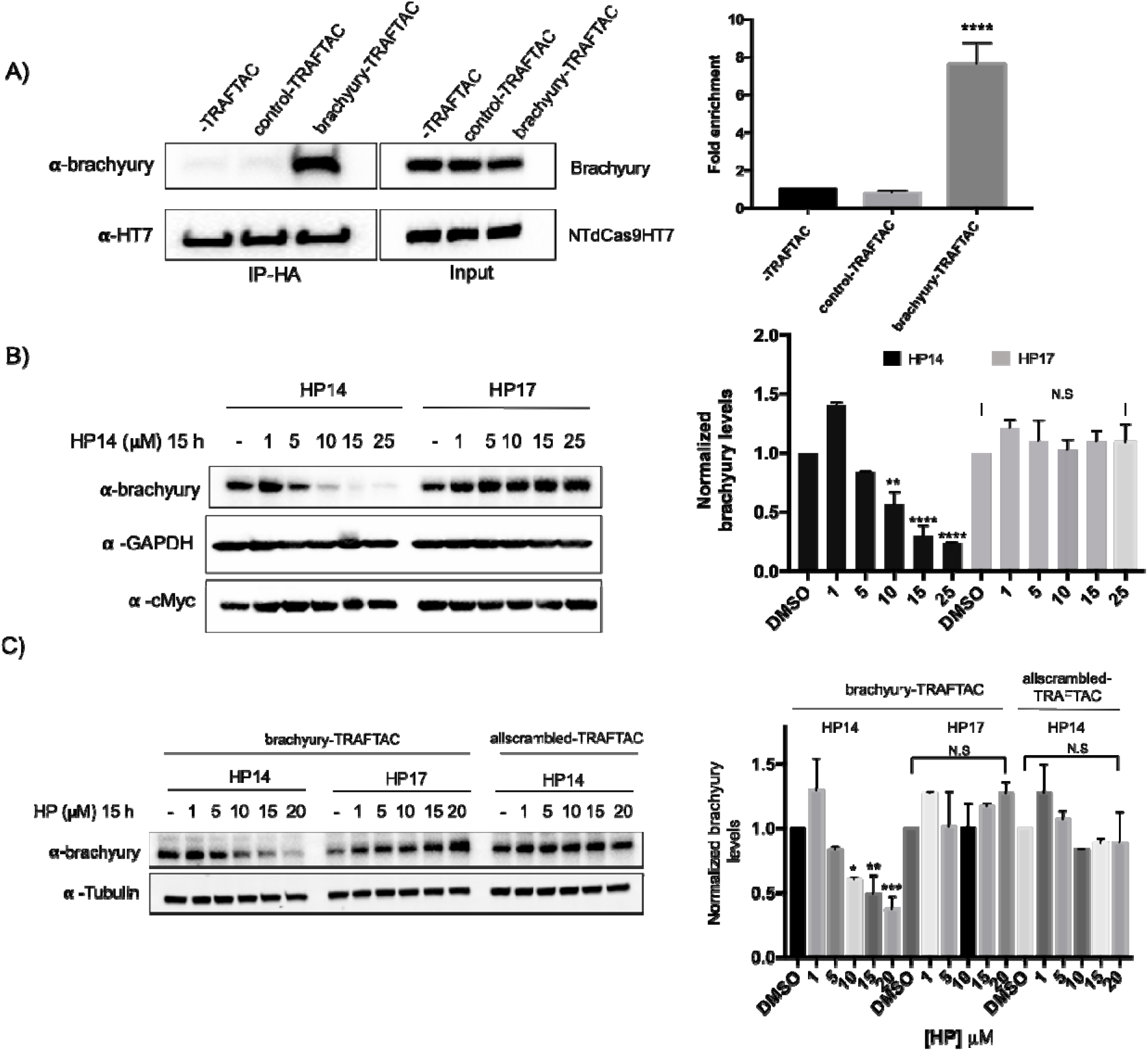
Endogenous brachyury degradation by TRAFTACs. A) Co-immunoprecipitation of brachyury in the presence and absence of brachyury-TRAFTAC or control-TRAFTAC oligos. Cell lysates were subjected to HA immunoprecipitation and eluted samples were probed for brachyury and HT7. B) Hela cells transiently transfected with NT-dCas9HT7 followed by a second transfection with brachyury-TRAFTAC. Then HP14 and HP17 were treated and cell lysates were prepared after 15 h and probed for brachyury, c-Myc and tubulin. C) Hela cells stably overexpressing NT-dCas9HT7 were transfected with either with brachyury-TRAFTAC or allscrambled-TRAFTAC followed by HP14 or HP17 treatment. Cell lysates were then probed for brachyury and tubulin levels. (Not significant (N.S); * P < 0.03; ** p<0.002; *** p<0.0002; **** p<0.0001)

## Discussion

Despite the many advancements in the field of chemotherapeutics, many cancer-driving proteins have remained intractable to current therapeutic modalities. Small molecule inhibitor development is one of the most accepted and clinically approved therapeutic strategies to treat many diseases including cancer. However, development of small molecule inhibitors, beyond those that target a few proteins classes (e.g. kinases and GPCRs) is challenging due to the lack of ligandable sites. Transcription factors are one such protein class that is difficult to drug. TFs regulate their downstream gene expression through a collective network of protein-protein and protein-DNA interactions, and lack enzymatic activity. Therefore, direct targeting of TFs is challenging. In this study, we have co-opted the DNA binding ability of TFs to develop TRAFTACs -- chimeric oligos that can simultaneously bind to a TOI and E3 ligase via an intermediary dCas9HT7 fusion protein. By employing TRAFTACs we have, for the first time, demonstrated that DNA-binding proteins, such as TFs can be degraded via the proteasomal pathway. We have successfully validated the proof-of-concept application of TRAFTAC technology in degrading transcription factors NF-κB and brachyury.

TNF-alpha induces IκB phosphorylation and its own subsequent degradation by the proteasome (independent of TRAFTAC). The dissociation of NF-κB from IκB inhibitory complex exposes the DNA binding region of NF-κB and facilitates its binding to the double-stranded DNA portion of the TRAFTAC chimeric oligo (52). Upon administration of the haloPROTAC, VHL E3 ligase recruitment induces subsequent proteasomal degradation of NF-κB. This feature of TRAFTACs offers an additional layer of selectivity to NF-κB proteasomal degradation. Hence, NF-κB-TRAFTACs could display a selectivity towards diseased cells with hyperactive NF-κB signaling, while sparing healthy cells with basal levels of NF-κB signaling (53). In TRAFTAC design we have adopted a HaloTag7 fusion of dCas9 protein as an intermediary protein. The dCas9HT7 fusion protein binds simultaneously to the TOI through the chimeric oligo and to the VHL-E3 complex through HP14. HaloPROTACs are efficient degraders of HaloTag fusion proteins but we have determined that the positioning of the HaloTag protein (N- or C-terminus) relative to dCas9HT7 significantly affected the resulting fusion protein’s susceptibility towards HP14-mediated degradation. While C-terminal HaloTag fusion is more prone to undergo HP14-mediated degradation at micromolar concentrations, the N-terminal HaloTag fusion is less susceptible. Our data also suggested that a potential use of C-terminal-HaloTag fusion Cas systems in CRISPR-mediated gene editing to achieve less off-target effects by inducing the degradation of engineered Cas proteins after the desired genome editing via administration of a haloPROTAC (HP3) (54–56).

Previous brachyury knockdown studies have demonstrated its critical roles in notochord fate during embryonic development. Although transgenic animal models such as knockout zebrafish models are available to study the biology of a given protein during embryonic development, the lack of spatiotemporal controllability of such systems hinders studying differential biology during different stages of embryonic development and/or post-developmentally. Here we have demonstrated that TRAFTACs can be used to induce conditional knockdown of brachyury in cells suggesting a potential future application in studying differential biology during embryonic development in animals when introduced to embryos as a ribonuclear complex. Also, TRAFTAC technology can be readily adapted to target other transcription factors by replacing the double-stranded DNA portion of the chimeric oligo. Thus, to our knowledge, TRAFTAC technology is the first degradation platform system that provides the flexibility to target many DNA-binding proteins for proteasomal degradation without devoting effort on TF ligand discovery.

Although many indirect strategies have been developed, only a few strategies have been demonstrated to directly inhibit transcription factor activity. Development of decoy elements that transiently inhibit TFs by interfering with its DNA binding ability has been successfully demonstrated (57–59). Further, decoys that directly bind at the minor grove of dsDNA have also been developed for many TFs including STAT3 and NF-κB (21, 60–62). However, these strategies function via *occupancy-driven* pharmacology, meaning these strategies provide only a transient blockade of the targeted transcription factors. To elicit robust inhibitory effects, these elements should display a sustained interaction with their target sites. Significantly, like PROTACs, TRAFTACs can exhibit an *event*-driven pharmacology that requires only transient interaction of TOI with the chimeric oligo to induce TF degradation. Furthermore, similarly to PROTACs, TRAFTACs also could be catalytic since the ternary complex of chimeric oligo:dCas9HT7:E3-ligase can bind to another molecule of the TOI and induce its degradation after having completed the first ubiquitination cycle of a previously-bound TOI. Therefore, transient interaction is enough to induce TOI degradation, in contrast current DNA-protein interaction inhibitors that possess limited inhibitory effects. Moreover, TRAFTACs are generalizable to many transcription factors with a known DNA binding sequence. We anticipate that most transcription factors could be successfully targeted for proteasomal degradation by changing the dsDNA portion of the chimeric oligo. Therefore, this method provides a straightforward strategy to target a broad range of DNA-binding proteins with minor changes to the system, thus offering an efficient approach to investigate unknown biology of known DNA-binding proteins in a rapid and robust way. Also, since TRAFTACs can be introduced as a ribonucleocomlplex, the current strategy would provide increased *in vivo* stability compared to short oligonucleotide derived-decoy elements.

As a future extension of this study, we anticipate modifying the current strategy to induce degradation of promoter-bound transcription factors in a gRNA-dependent and locus-dependent manner. This strategy would also provide an avenue to repress a single, targeted gene by inducing the degradation of its promoter-bound transcription factor without altering the expression of other genes that are controlled by the same transcription factor. Collectively, the TRAFTAC technology holds great potential, both as a chemical biology tool and a therapeutic strategy for several reasons: 1) many oncogenic transcription factors have already been identified as potential drug targets; 2) synthesis of the chimeric oligo is simple and straightforward, 3) chimeric oligos are readily adaptable to target different TFs; 4) structural information of the transcription factor is not necessary; 5) laborious small molecule ligand discovery campaigns are not required, and finally; 6) TRAFTACs only require transient interaction with the target transcription factor to induce TF degradation relative to decoy or antisense oligonucleotide approaches, which require persistent engagement with target proteins to achieve the desired inhibitory effects. In fact, TRAFTAC could be a potential therapeutic strategy to treat any disease in which a transcription factor activity is dysregulated. Once coupled with an efficient delivery strategy, TRAFTAC technology holds tremendous potential to target hard-to-drug transcription factors and other DNA-binding proteins for proteasomal degradation in patients in the future.

## Acknowledgement

We thank Dr. John Hines and Dr. Dhanusha A. Nalawansha for the critical reading of the manuscript, insightful comments and critiques. C.M.C. is funded by the NIH (R35CA197589) and is supported by an American Cancer Research Professorship. (Figures were created with BioRender.com)

## Reference

1. K. M. Sakamoto et al., Protacs: Chimeric molecules that target proteins to the Skp1–Cullin–F box complex for ubiquitination and degradation. Proceedings of the National Academy of Sciences 98, 8554 (2001).

2. A. C. Lai, C. M. Crews, Induced protein degradation: an emerging drug discovery paradigm. Nature Reviews Drug Discovery 16, 101–114 (2017).

3. G. Nalepa, M. Rolfe, J. W. Harper, Drug discovery in the ubiquitin–proteasome system. Nature Reviews Drug Discovery 5, 596–613 (2006).

4. H. Zhang et al., Discovery of potent epidermal growth factor receptor (EGFR) degraders by proteolysis targeting chimera (PROTAC). European Journal of Medicinal Chemistry 189, 112061 (2020).

5. A. C. Lai et al., Modular PROTAC Design for the Degradation of Oncogenic BCR-ABL. Angew Chem Int Ed Engl 55, 807–810 (2016).

6. G. M. Burslem et al., Targeting BCR-ABL1 in Chronic Myeloid Leukemia by PROTAC-Mediated Targeted Protein Degradation. Cancer research 79, 4744–4753 (2019).

7. G. M. Burslem, J. Song, X. Chen, J. Hines, C. M. Crews, Enhancing Antiproliferative Activity and Selectivity of a FLT-3 Inhibitor by Proteolysis Targeting Chimera Conversion. Journal of the American Chemical Society 140, 16428–16432 (2018).

8. X. Zhang, V. M. Crowley, T. G. Wucherpfennig, M. M. Dix, B. F. Cravatt, Electrophilic PROTACs that degrade nuclear proteins by engaging DCAF16. Nature Chemical Biology 15, 737–746 (2019).

9. G. L. Verdine, L. D. Walensky, The challenge of drugging undruggable targets in cancer: lessons learned from targeting BCL-2 family members. Clinical cancer research: an official journal of the American Association for Cancer Research 13, 7264–7270 (2007).

10. J. S. Lazo, E. R. Sharlow, Drugging Undruggable Molecular Cancer Targets. Annual Review of Pharmacology and Toxicology 56, 23–40 (2016).

11. T. Sharifnia et al., Small-molecule targeting of brachyury transcription factor addiction in chordoma. Nature Medicine 25, 292–300 (2019).

12. S. Ali et al., Molecular mechanisms and mode of tamoxifen resistance in breast cancer. Bioinformation 12, 135–139 (2016).

13. C. K. Osborne, A. Wakeling, R. I. Nicholson, Fulvestrant: an oestrogen receptor antagonist with a novel mechanism of action. British journal of cancer 90 Suppl 1, S2–6 (2004).

14. M. A. Rice, S. V. Malhotra, T. Stoyanova, Second-Generation Antiandrogens: From Discovery to Standard of Care in Castration Resistant Prostate Cancer. Front Oncol 9, 801–801 (2019).

15. P. A. Watson, V. K. Arora, C. L. Sawyers, Emerging mechanisms of resistance to androgen receptor inhibitors in prostate cancer. Nature Reviews Cancer 15, 701–711 (2015).

16. J. Salami et al., Androgen receptor degradation by the proteolysis-targeting chimera ARCC-4 outperforms enzalutamide in cellular models of prostate cancer drug resistance. Communications Biology 1, 100 (2018).

17. J. Hu et al., Discovery of ERD-308 as a Highly Potent Proteolysis Targeting Chimera (PROTAC) Degrader of Estrogen Receptor (ER). Journal of medicinal chemistry 62, 1420–1442 (2019).

18. X. Sun et al., PROTACs: great opportunities for academia and industry. Signal Transduction and Targeted Therapy 4, 64 (2019).

19. L. Bai et al., A Potent and Selective Small-Molecule Degrader of STAT3 Achieves Complete Tumor Regression <em>In Vivo</em>. Cancer Cell 36, 498–511.e417 (2019).

20. F. Yang et al., Antitumor activity of a pyrrole-imidazole polyamide. Proceedings of the National Academy of Sciences 110, 1863 (2013).

21. N. R. Wurtz, J. L. Pomerantz, D. Baltimore, P. B. Dervan, Inhibition of DNA Binding by NF-κB with Pyrrole-Imidazole Polyamides. Biochemistry 41, 7604–7609 (2002).

22. A. J. Walhout, J. M. Gubbels, R. Bernards, P. C. van der Vliet, H. T. Timmers, c-Myc/Max heterodimers bind cooperatively to the E-box sequences located in the first intron of the rat ornithine decarboxylase (ODC) gene. Nucleic Acids Res 25, 1493–1501 (1997).

23. P. Raffeiner et al., In vivo quantification and perturbation of Myc-Max interactions and the impact on oncogenic potential. Oncotarget 5, 8869–8878 (2014).

24. A. Castell et al., A selective high affinity MYC-binding compound inhibits MYC:MAX interaction and MYC-dependent tumor cell proliferation. Scientific Reports 8, 10064 (2018).

25. S. H. Choi et al., Targeted Disruption of Myc–Max Oncoprotein Complex by a Small Molecule. ACS Chemical Biology 12, 2715–2719 (2017).

26. H. Jiang et al., Stabilizers of the Max homodimer identified in virtual ligand screening inhibit Myc function. Mol Pharmacol 76, 491–502 (2009).

27. N. B. Struntz et al., Stabilization of the Max Homodimer with a Small Molecule Attenuates Myc-Driven Transcription. Cell chemical biology 26, 711–723.e714 (2019).

28. Y. Chen et al., Bt354 as a new STAT3 signaling pathway inhibitor against triple negative breast cancer. Journal of drug targeting 26, 920–930 (2018).

29. X. Dai et al., Osthole inhibits triple negative breast cancer cells by suppressing STAT3. Journal of experimental & clinical cancer research: CR 37, 322 (2018).

30. C. Y. Liu et al., The tyrosine kinase inhibitor nintedanib activates SHP-1 and induces apoptosis in triple-negative breast cancer cells. Experimental & molecular medicine 49, e366 (2017).

31. T. J. Rios-Fuller et al., Ganoderma lucidum extract (GLE) impairs breast cancer stem cells by targeting the STAT3 pathway. Oncotarget 9, 35907–35921 (2018).

32. S. A. Reddy, J. H. Huang, W. S. Liao, Phosphatidylinositol 3-kinase in interleukin 1 signaling. Physical interaction with the interleukin 1 receptor and requirement in NFkappaB and AP-1 activation. The Journal of biological chemistry 272, 29167–29173 (1997).

33. M. Karin, Y. Yamamoto, Q. M. Wang, The IKK NF-kappa B system: a treasure trove for drug development. Nature reviews. Drug discovery 3, 17–26 (2004).

34. V. Pande, M. J. Ramos, NF-kappaB in human disease: current inhibitors and prospects for de novo structure based design of inhibitors. Current medicinal chemistry 12, 357–374 (2005).

35. K. Raina et al., PROTAC-induced BET protein degradation as a therapy for castration-resistant prostate cancer. Proceedings of the National Academy of Sciences 113, 7124 (2016).

36. G. E. Winter et al., Phthalimide conjugation as a strategy for in vivo target protein degradation. Science 10.1126/science.aab1433, aab1433 (2015).

37. F. E. Chen, D. B. Huang, Y. Q. Chen, G. Ghosh, Crystal structure of p50/p65 heterodimer of transcription factor NF-kappaB bound to DNA. Nature 391, 410–413 (1998).

38. L. S. Qi et al., Repurposing CRISPR as an RNA-guided platform for sequencespecific control of gene expression. Cell 152, 1173–1183 (2013).

39. D. L. Buckley et al., HaloPROTACS: Use of Small Molecule PROTACs to Induce Degradation of HaloTag Fusion Proteins. ACS chemical biology 10, 1831–1837 (2015).

40. K. H. Mellits, R. T. Hay, S. Goodbourn, Proteolytic degradation of MAD3 (I kappa B alpha) and enhanced processing of the NF-kappa B precursor p105 are obligatory steps in the activation of NF-kappa B. Nucleic Acids Res 21, 5059–5066 (1993).

41. B. E. Smith et al., Differential PROTAC substrate specificity dictated by orientation of recruited E3 ligase. Nature Communications 10, 131 (2019).

42. A. D. Buhimschi et al., Targeting the C481S Ibrutinib-Resistance Mutation in Bruton’s Tyrosine Kinase Using PROTAC-Mediated Degradation. Biochemistry 57, 3564–3575 (2018).

43. S. Jaime-Figueroa, A. D. Buhimschi, M. Toure, J. Hines, C. M. Crews, Design, synthesis and biological evaluation of Proteolysis Targeting Chimeras (PROTACs) as a BTK degraders with improved pharmacokinetic properties. Bioorganic & Medicinal Chemistry Letters 30, 126877 (2020).

44. H. Nishimasu et al., Crystal Structure of Cas9 in Complex with Guide RNA and Target DNA. Cell 156, 935–949 (2014).

45. J. Zhu, K. M. Kwan, S. Mackem, Putative oncogene &lt;em&gt;Brachyury&lt;/em&gt; (&lt;em&gt;T&lt;/em&gt;) is essential to specify cell fate but dispensable for notochord progenitor proliferation and EMT. Proceedings of the National Academy of Sciences 113, 3820 (2016).

46. D. L. Stemple et al., Mutations affecting development of the notochord in zebrafish. Development 123, 117 (1996).

47. M. Chen et al., The Roles of Embryonic Transcription Factor BRACHYURY in Tumorigenesis and Progression. Front Oncol 10 (2020).

48. R. Bosotti et al., Establishment and genomic characterization of the new chordoma cell line Chor-IN-1. Scientific reports 7, 9226–9226 (2017).

49. N. Presneau et al., Role of the transcription factor T (brachyury) in the pathogenesis of sporadic chordoma: a genetic and functional-based study. The Journal of pathology 223, 327–335 (2011).

50. X. R. Yang et al., T (brachyury) gene duplication confers major susceptibility to familial chordoma. Nature genetics 41, 1176–1178 (2009).

51. N. Pillay et al., A common single-nucleotide variant in T is strongly associated with chordoma. Nature genetics 44, 1185–1187 (2012).

52. M. S. Hayden, S. Ghosh, Regulation of NF-κB by TNF family cytokines. Semin Immunol 26, 253–266 (2014).

53. L. Xia et al., Role of the NFκB-signaling pathway in cancer. Onco Targets Ther 11, 2063–2073 (2018).

54. S. W. Cho et al., Analysis of off-target effects of CRISPR/Cas-derived RNA-guided endonucleases and nickases. Genome Res 24, 132–141 (2014).

55. S. A. Gangopadhyay et al., Precision Control of CRISPR-Cas9 Using Small Molecules and Light. Biochemistry 58, 234–244 (2019).

56. Z. Tu et al., Promoting Cas9 degradation reduces mosaic mutations in nonhuman primate embryos. Sci Rep 7, 42081 (2017).

57. R. Morishita, N. Tomita, Y. Kaneda, T. Ogihara, Molecular therapy to inhibit NFkappaB activation by transcription factor decoy oligonucleotides. Current opinion in pharmacology 4, 139–146 (2004).

58. D. De Stefano, Oligonucleotides decoy to NF-kappaB: becoming a reality? Discovery medicine 12, 97–105 (2011).

59. K. Egashira et al., Long-term follow up of initial clinical cases with NF-kappaB decoy oligodeoxynucleotide transfection at the site of coronary stenting. The journal of gene medicine 10, 805–809 (2008).

60. M. Sen et al., First-in-human trial of a STAT3 decoy oligonucleotide in head and neck tumors: implications for cancer therapy. Cancer Discov 2, 694–705 (2012).

61. D. S. Lee, R. A. O’Keefe, P. K. Ha, J. R. Grandis, D. E. Johnson, Biochemical Properties of a Decoy Oligodeoxynucleotide Inhibitor of STAT3 Transcription Factor. Int J Mol Sci 19, 1608 (2018).

62. S. Imbaby et al., Beneficial effect of STAT3 decoy oligodeoxynucleotide transfection on organ injury and mortality in mice with cecal ligation and puncture-induced sepsis. Scientific Reports 10, 15316 (2020).

