## Supplemental Materials for "Targeted Degradation of Transcription Factors by TRAFTACs: Transcription Factor Targeting Chimeras"

**EXPERIMENTAL SECTION**

**Chemical syntheses**

**SJF-7432 = HP13**

**SJF-7434 = HP14**

**SJF-4625 = HP15**

**SJF-4627 = HP16**

**JH-6073	= HP17**

**General comments.** Unless otherwise indicated, common reagents or materials were obtained from commercial source and used without further purification. Tetrahydrofuran (THF), dimethylformamide (DMF), and Dichloromethane (CH_2_Cl_2_) were dried by a PureSolv^TM^ solvent drying system. Flash column chromatography was performed using silica gel 60 (230-400 mesh). Analytical (TLC) and preparative (PTLC) thin layer chromatography was carried out on Merck silica gel plates with QF-254 indicator and visualized by UV or iodine. ^1^H and ^13^C NMR spectra were recorded on an Agilent DD_2_ 500 (500 MHz ^1^H; 125 MHz ^13^C) or Agilent DD_2_ 600 (600 MHz ^1^H; 150 MHz ^13^C) or Agilent DD_2_ 400 (400 MHz ^1^H; 100 MHz ^13^C) spectrometer at room temperature. Chemical shifts were reported in ppm relative to the residual CDCl_3_ (δ 7.26 ppm ^1^H; δ 77.00 ppm ^13^C), CD_3_OD (δ 3.31 ppm ^1^H; δ 49.00 ppm ^13^C), or *d^6^*-DMSO (δ 2.50 ppm ^1^H; δ 39.52 ppm ^13^C). NMR chemical shifts were expressed in ppm relative to internal solvent peaks, and coupling constants were measured in Hz. (bs = broad signal). Mass spectra were obtained using electrospray ionization (ESI) on a time of flight (TOF) mass spectrometer. **VHL** ligands **7** and **10** were prepared according with the literature^1^ or acquired commercially.

**Scheme 1.- Synthetic Approach for SJF-7432, SJF-7434, SJF-4625 and JH-6073**

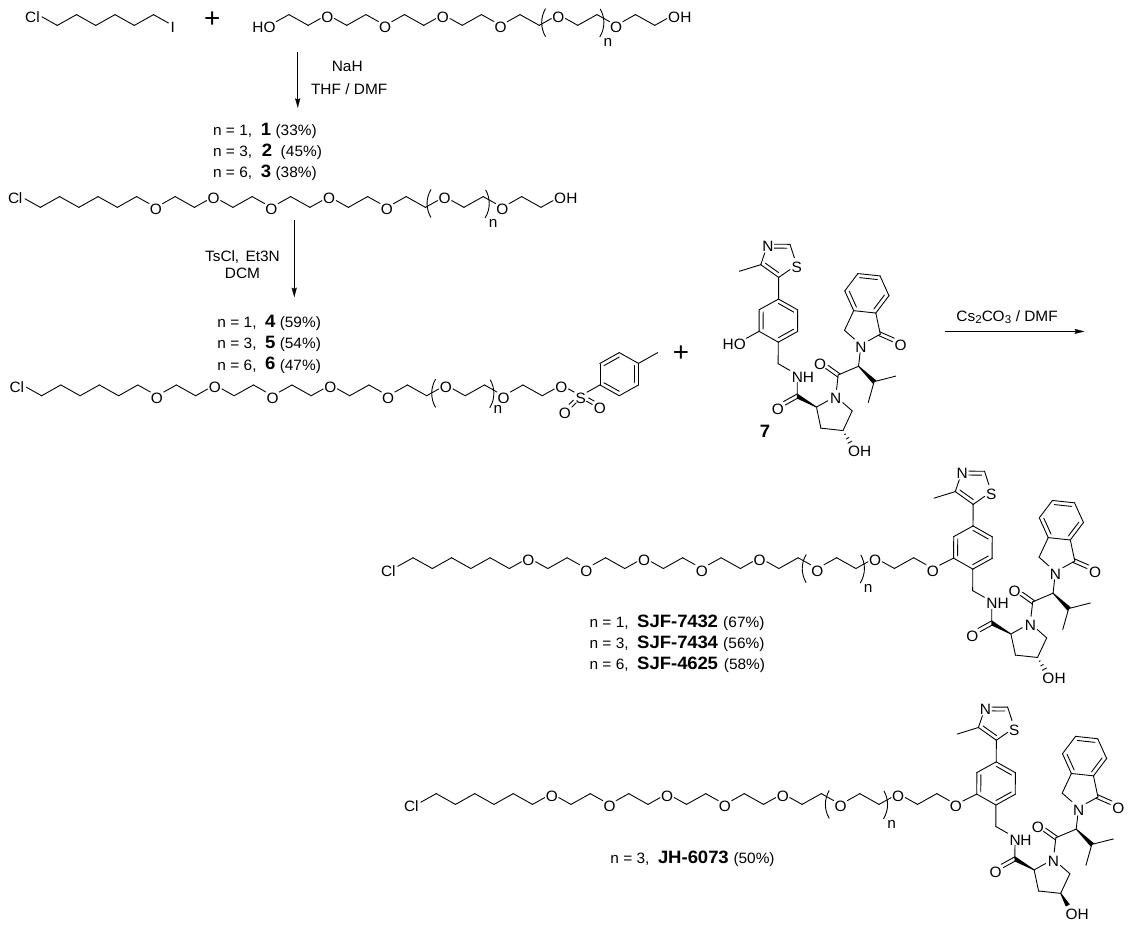

**Scheme 2.- Synthetic Approach for SJF-4627**

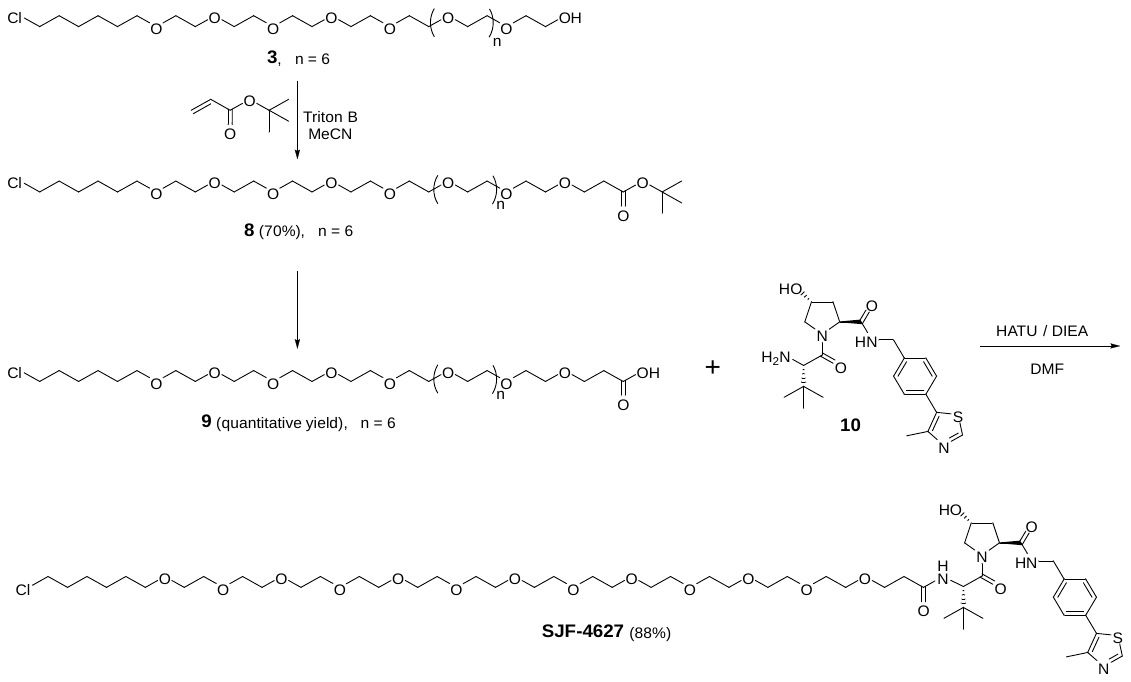

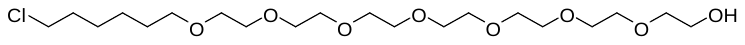

**27-chloro-3,6,9,12,15,18,21-heptaoxaheptacosan-1-ol (1).** To a solution of 3,6,9,12,15,18-hexaoxaicosane-1,20-diol (1.06 mL, 3.65 mmol) in a mixture of DMF (5 mL ) and THF (5 mL) was added NaH (60.0 %, 81.1 mg, 2.03 mmol) at room temperature under argon. After 40 minutes, 1-chloro-6-iodo-hexane (0.123 mL, 0.811 mmol) was added, and the mixture was stirred at room temperature for 16 h (overnight). The mixture was then quenched with diluted with 1 M HCl (50 mL) and extracted with Ethyl acetate (50 mL). The organic layer was dried over Na_2_SO_4_ and the solvent removed under reduced pressure. The crude product was purified by column chromatography (Gradient, DCM 100% to DCM:MeOH, 9:1) to give 120 mg of product as an oil (33% yield). ^1^H NMR (500 MHz, Chloroform-d) δ 3.74 – 3.69 (m, 2H), 3.69 – 3.58 (m, 26H), 3.57 (t, J = 5.3 Hz, 2H), 3.44 (t, J = 6.6 Hz, 2H), 2.55 (bs, 1H), 1.81 – 1.71 (m, 2H), 1.58 (p, J = 6.8 Hz, 3H), 1.50 – 1.30 (m, 4H). ^13^C NMR (151 MHz, Chloroform-d) δ 72.59, 71.35, 70.71, 70.71, 70.70, 70.69, 70.68, 70.67, 70.67, 70.66, 70.64, 70.44, 70.43, 70.21, 61.83, 45.18, 32.67, 29.58, 26.83, 25.55. LC-MS (ESI); m/z [M+1]^+^; Calcd. C_20_H_42_ClO_8_, 445.2568. Found 445.38.

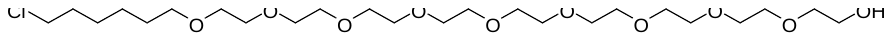

**33-chloro-3,6,9,12,15,18,21,24,27-nonaoxatritriacontan-1-ol** (**2**). To a solution of 3,6,9,12,15,18,21,24-octaoxahexacosane-1,26-diol (1.68 mL, 4.56 mmol) in a mixture of DMF (5 mL ) and THF (5 mL) was added NaH (60.0 %, 150 mg, 3.75 mmol) at room temperature under argon. After 40 minutes, 1-chloro-6-iodo-hexane (0.154 mL, 1.01 mmol) was added, and the mixture was stirred at room temperature for 16 h (overnight). The mixture was then quenched with a mixture of diluted 1 M HCl (5 mL) and brine (15 mL), and then the reaction mixture was extracted with Ethyl acetate (50 mL). The organic layer was dried over Na_2_SO_4_ and the solvent removed under reduced pressure. The crude product was purified by column chromatography (Gradient, DCM 100% to DCM:MeOH, 9:1) to give 242 mg of pure product (45% yield).. ^1^H NMR (500 MHz, Chloroform-d) δ 3.84 – 3.54 (m, 36H), 3.51 (t, J = 7.1 Hz, 2H), 3.44 (t, J = 6.6 Hz, 2H), 2.76 (bs, 1H), 1.76 (p, J = 7.0 Hz, 2H), 1.58 (p, J = 6.8 Hz, 2H), 1.48 – 1.30 (m, 4H). ^13^C NMR (151 MHz, Chloroform-d) δ 72.66, 71.34, 70.72, 70.70, 70.66 (9C), 70.64, 70.62, 70.62, 70.40, 70.21, 61.81, 45.17, 32.66, 29.57, 26.82, 25.54. LC-MS (ESI); m/z [M+1]^+^; Calcd. C_24_H_50_ClO_10_, 533.3092. Found 533.45.

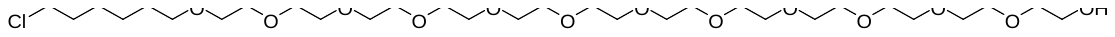

**42-chloro-3,6,9,12,15,18,21,24,27,30,33,36-dodecaoxadotetracontan-1-ol** (**3**). To a solution of 3,6,9,12,15,18,21,24,27,30,33-undecaoxapentatriacontane-1,35-diol (998 mg, 1.83 mmol) in a mixture of DMF (3 mL ) and THF (3 mL) was added NaH (60.0 %, 48.7 mg, 1.22 mmol) at room temperature under argon. After 40 minutes, 1-chloro-6-iodo-hexane (0.0616 mL, 0.406 mmol) was added, and the mixture was stirred at room temperature for 16 h (overnight). The mixture was then quenched with a mixture of diluted 1 M HCl (5 mL) and brine (20 mL), and then the reaction mixture was extracted with Ethyl acetate (50 mL). The organic layer was dried over Na_2_SO_4_ and the solvent removed under reduced pressure. The crude product was purified by column chromatography (Gradient, DCM 100% to DCM:MeOH, 7:3) to give 104 mg of pure product as an oil (38% yield) ^1^H (400 MHz, Chloroform-d) δ 3.93 – 3.55 (m, 48H), 3.51 (t, J = 6.7 Hz, 2H), 3.44 (t, J = 6.6 Hz, 2H), 2.56 (bs, 1H), 1.76 (pd, J = 6.7, 2.1 Hz, 2H), 1.64 – 1.50 (m, 2H), 1.50 – 1.27 (m, 4H). ^13^C NMR (151 MHz, cdcl3) δ 72.63, 71.34, 70.72, 70.70, 70.68, 70.66 (15C), 70.63, 70.42, 70.42, 70.21, 61.82, 45.17, 32.65, 29.56, 26.81, 25.53. LC-MS (ESI); m/z [M+1]^+^; Calcd. C_30_H_62_ClO_13_, 665.3878. Found 665.54.

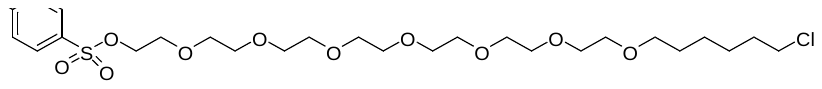

**27-chloro-3,6,9,12,15,18,21-heptaoxaheptacosyl 4-methylbenzenesulfonate** (**4**). To a solution of 27-chloro-3,6,9,12,15,18,21-heptaoxaheptacosan-1-ol (114 mg, 0.256 mmol) and TEA (0.214 mL, 1.54 mmol) in DCM (5 ml) was added toluene-4-sulfonyl chloride (63.5 mg, 0.333 mmol) at room temperature. The reaction mixture was stirred for 5 at the same temperature. Then the reaction mixture was poured into aqueous NaHCO_3_ (sat. solution, 10 mL) and product was extracted with DCM (2x20 mL). Organic extracts were combined, dried (Na_2_SO_4_) and evaporated under vacuum. Crude product was purified by column chromatography (DCM 100% to DCM:MeOH, 95:5), to give 90 mg (58% yield) of product as an oil. ^1^H NMR (500 MHz, Chloroform-d) δ 7.79 (d, J = 10.1 Hz, 2H), 7.34 (d, J = 6.6 Hz, 2H), 4.15 (t, J = 6.1 Hz, 2H), 3.81 – 3.55 (m, 24H), 3.53 (t, J = 6.7 Hz, 2H), 3.45 (t, J = 7.9 Hz, 2H), 2.44 (s, 3H), 1.77 (p, J = 9.2 Hz, 2H), 1.59 (p, J = 7.8, 6.3 Hz, 2H), 1.52 – 1.22 (m, 4H). ^13^C NMR (151 MHz, Chloroform-d) δ 144.89, 133.13, 129.93, 128.10, 71.35, 70.87, 70.73, 70.71, 70.69 (6C), 70.68, 70.64, 70.22, 69.36, 68.80, 45.18, 32.67, 29.58, 26.82, 25.55, 21.77. LC-MS (ESI); m/z [M+1]^+^ ; calcd. C_27_H_48_ClO_10_S, 599.2656. Found 599.37.

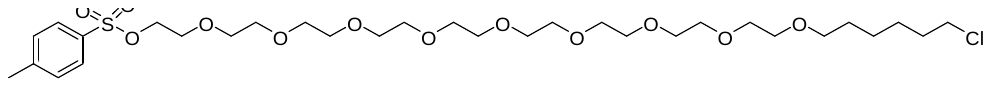

**33-chloro-3,6,9,12,15,18,21,24,27-nonaoxatritriacontyl 4-methylbenzenesulfonate** (**5**). To a solution of 33-chloro-3,6,9,12,15,18,21,24,27-nonaoxatritriacontan-1-ol (230 mg, 0.431 mmol) and TEA (0.361 mL, 2.59 mmol) in DCM (5 ml) was added Toluene-4-sulfonyl chloride (107 mg, 0.561 mmol) at room temperature. The reaction mixture was stirred for 5 at the same temperature. Then the reaction mixture was poured into aqueous NaHCO_3_ (sat. solution, 10 mL) and product was extracted with DCM (2x20 mL). Organic extracts were combined, dried (Na_2_SO_4_) and evaporated under vacuum. Crude product was purified by column chromatography (DCM 100% to DCM:MeOH, 95:5), to give 161 mg (54% yield) of product. ^1^H NMR (600 MHz, DMSO-d6) δ 7.78 (d, J = 7.6 Hz, 2H), 7.48 (d, J = 7.7 Hz, 2H), 4.11 (t, J = 5.2 Hz, 2H), 3.61 (t, J = 6.3 Hz, 2H), 3.57 (t, J = 4.3 Hz, 2H), 3.47 (d, J = 32.9 Hz, 32H), 3.36 (t, J = 6.1 Hz, 2H), 2.42 (s, 3H), 1.70 (p, J = 6.7 Hz, 2H), 1.48 (p, J = 6.6 Hz, 2H), 1.38 (p, J = 7.1 Hz, 2H), 1.30 (p, J = 7.5 Hz, 2H). ^13^C NMR (151 MHz, DMSO-d6) δ 144.86, 132.41, 130.11, 127.61, 70.16, 69.97, 69.81, 69.78 (6C), 69.75, 69.70, 69.65, 69.48, 67.88, 45.35, 32.02, 29.05, 26.11, 24.93, 21.08. LC-MS (ESI); m/z [M+1]^+^; Calcd. C_31_H_56_ClO_12_S, 687.3181. Found 687.44.

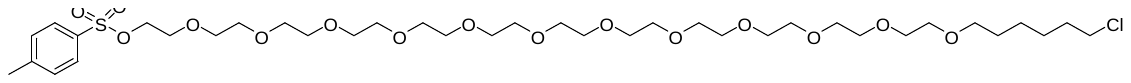

**42-chloro-3,6,9,12,15,18,21,24,27,30,33,36-dodecaoxadotetracontyl 4-methylbenzenesulfonate** (**6**). To a solution of 42-chloro-3,6,9,12,15,18,21,24,27,30,33,36-dodecaoxadotetracontan-1-ol (48.0 mg, 0.0722 mmol) and TEA (0.0603 mL, 0.433 mmol) in DCM (2 ml) was added Toluene-4-sulfonyl chloride (17.9 mg, 0.0938 mmol) at room temperature. The reaction mixture was stirred for 5h at the same temperature. Then the reaction mixture was poured into aqueous NaHCO_3_ (sat. solution, 10 mL) and product was extracted with DCM (2x20 mL). Organic extracts were combined, dried (Na_2_SO_4_) and evaporated under vacuum. Crude product was purified by PTLC (DCM:MeOH:NH_4_OH, 90:9:1), to give 28mg (47% yield) of pure product. ^1^H NMR (400 MHz, DMSO-d6) δ 7.78 (d, J = 8.3 Hz, 1H), 7.48 (d, J = 8.0 Hz, 2H), 4.23 – 3.99 (m, 2H), 3.61 (t, J = 6.6 Hz, 2H), 3.59 – 3.40 (m, 46H), 3.36 (t, J = 6.5 Hz, 2H), 2.42 (s, 3H), 1.77 – 1.62 (m, 2H), 1.48 (p, J = 6.8 Hz, 2H), 1.43 – 1.17 (m, 4H). ^13^C NMR (151 MHz, dmso) δ 144.90, 132.42, 130.14, 127.64, 70.18, 69.99, 69.83, 69.79 (C18), 69.71, 69.67, 69.51, 67.89, 45.37, 32.04, 29.07, 26.13, 24.95, 21.10. LC-MS (ESI); m/z [M+Na]^+^; Calcd. C_37_H_67_ClO_15_SNa, 841.3787.

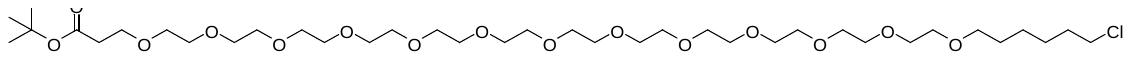

**tert-butyl 46-chloro-4,7,10,13,16,19,22,25,28,31,34,37,40-tridecaoxahexatetracontanoate** (**8**). To a solution of 42-chloro-3,6,9,12,15,18,21,24,27,30,33,36-dodecaoxadotetracontan-1-ol (**3**) (50.0 mg, 0.0752 mmol) in acetonitrile (2 mL) was added tert-butyl prop-2-enoate (0.218 mL, 1.50 mmol) followed by  Triton B (40.0 %, 0.158 mL, 0.400 mmol, in 40% by weight in water). The mixture was stirred at room temperature for 48 hours. The mixture was concentrated under vacuum and crude product was purified by PTLC (DCM:MeOH:NH_4_OH, 90:9:1) to give 42 mg of product as an oil (70% yield). ^1^H NMR (400 MHz, Chloroform-d) δ 3.68 (t, J = 6.6 Hz, 2H), 3.65 – 3.54 (m, 48H), 3.51 (t, J = 6.7 Hz, 2H), 3.43 (t, J = 6.6 Hz, 2H), 2.48 (t, J = 6.6 Hz, 2H), 1.75 (p, J = 13.5, 6.2 Hz, 2H), 1.64 – 1.50 (m, 2H), 1.42 (s, 9H), 1.50 – 1.26 (m, 4H). ^13^C NMR (151 MHz, dmso) δ 170.42, 79.71, 70.19, 69.84, 69.81(20C), 69.72, 69.69, 69.52, 66.24, 45.38, 35.85, 32.04, 29.08, 27.75, 26.13, 24.95. LC-MS (ESI); m/z [M+Na]^+^: Calcd. for C_37_H_73_ClO_15_Na, 815.4535. Found 815.4527.

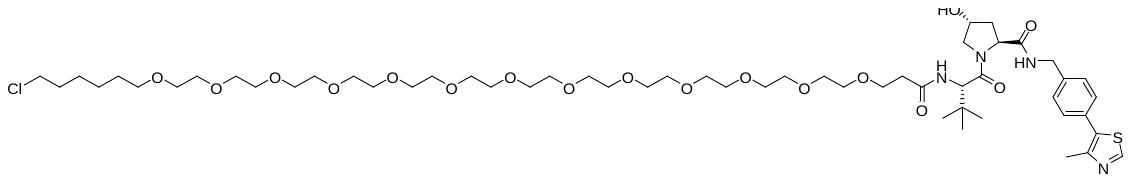

**(2S,4R)-1-((S)-2-(tert-butyl)-49-chloro-4-oxo-7,10,13,16,19,22,25,28,31,34,37,40,43-tridecaoxa-3-azanonatetracontanoyl)-4-hydroxy-N-(4-(4-methylthiazol-5-yl)benzyl)pyrrolidine-2-carboxamide** (**SJF-4627**). A solution of tert-butyl 46-chloro-4,7,10,13,16,19,22,25,28,31,34,37,40-tridecaoxahexatetra contanoate (**8**) (18.0 mg, 0.0227 mmol) in a mixture of TFA (1 ml, 13.46 mmol) and Dichloromethane (2 ml) was stirred for 2 h. Then the solvent was removed under vacuum and crude product was dried under high vacuum for 2 h. Crude product (**9**) was used in the next step without any further purification (16.7 mg, quantitative yield). HRMS (ESI); m/z: [M+H]^+^ Calcd. for C_33_H_66_ClO_15_, 737.4090. Found 737.4090.

To a solution of 46-chloro-4,7,10,13,16,19,22,25,28,31,34,37,40-tridecaoxahexatetracontanoic acid (16.7 mg, 0.0226 mmol) and (2S,4R)-1-[(2S)-2-amino-3,3-dimethyl-butanoyl]-4-hydroxy-N-[[4-(4-methylthiazol-5-yl)phenyl]methyl]pyrrolidine-2-carboxamide;hydrochloride (12.7 mg, 0.0272 mmol) in DMF (2 ml) was added N,N-Diisopropylethylamine (0.298 mL, 1.71 mmol) and O-(7-Azabenzotriazol-1-yl)-N,N,N',N'-tetramethyluronium hexafluorophosphate (12.9 mg, 0.0340 mmol) at room temperature. The reaction mixture was stirred for 12 h (overnight) at the same temperature. TLC (DCM:MB, 1:1) shows no starting materials. Reaction mixture was diluted with EtOAc (10 mL), washed with water (4x10 mL), dried (Na_2_SO_4_) and evaporated under vacuum. Crude product was purified by PTLC (DCM:MeOH:NH_4_OH, 90:9:1), to 23 mg of product (88 % yield). ^1^H NMR (500 MHz, DMSO-d6) δ 8.98 (s, 1H), 8.56 (t, J = 5.9 Hz, 1H), 7.91 (d, J = 9.3 Hz, 1H), 7.42 (d, J = 8.2 Hz, 2H), 7.38 (d, J = 8.2 Hz, 2H), 5.12 (d, J = 3.4 Hz, 1H), 4.55 (d, J = 9.4 Hz, 1H), 4.46 – 4.39 (m, 2H), 4.37 – 4.32 (m, 1H), 4.22 (dd, J = 15.8, 5.2 Hz, 1H), 4.00 – 3.42 (m, 54H), 3.37 (t, J = 6.5 Hz, 2H), 2.58 – 2.51 (m, 1H), 2.44 (s, 3H), 2.39 – 2.32 (m, 1H), 2.07 – 2.00 (m, 1H), 1.95 – 1.87 (m, 1H), 1.70 (dt, J = 14.3, 6.7 Hz, 2H), 1.48 (p, J = 13.8, 6.7 Hz, 2H), 1.42 – 1.25 (m, 4H), 0.94 (s, 9H). ^13^C NMR (151 MHz, dmso) δ 171.96, 169.96, 169.54, 151.48, 147.73, 139.53, 131.18, 129.65, 128.65, 127.44, 70.19, 69.84, 69.80, 69.73, 69.52, 69.50, 68.89 (20C), 66.97, 58.73, 56.40, 56.31, 45.39, 41.66, 37.96, 35.67, 35.39, 32.04, 29.08, 26.34, 26.13, 24.95, 15.96. HRMS (ESI); m/z [M+H]^+^: Calcd. for C_55_H_94_ClN_4_O_17_S, 1149.6023. Found 1149.6023.

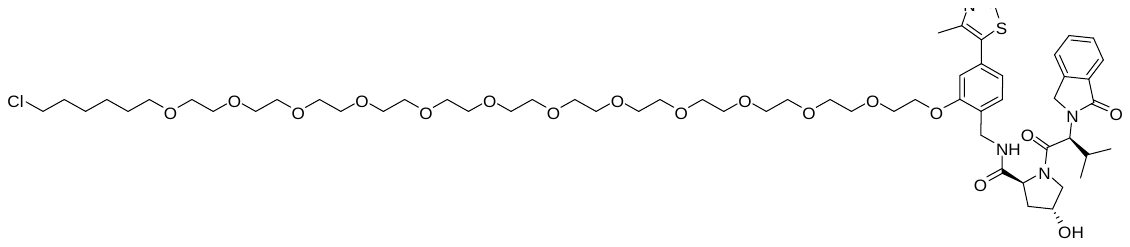

**(2S,4R)-N-(2-((42-chloro-3,6,9,12,15,18,21,24,27,30,33,36-dodecaoxadotetracontyl)oxy)-4-(4-methyl -thiazol-5-yl)benzyl)-4-hydroxy-1-((S)-3-methyl-2-(1-oxoisoindolin-2-yl)butanoyl)pyrrolidine-2-carboxa -mide** (**SJF-4625**). To a mixture of (2S,4R)-4-hydroxy-N-[[2-hydroxy-4-(4-methylthiazol-5-yl)phenyl]methyl]-1-[(2S)-3-methyl-2-(1-oxoisoindolin-2-yl)butanoyl]pyrrolidine-2-carboxamide (22.0 mg, 0.0402 mmol) and 27-chloro-3,6,9,12,15,18,21-heptaoxaheptacosyl 4-methylbenzenesulfonate (**6**) (23.0 mg, 0.0335 mmol) in DMF (1 mL) was added Cs_2_CO_3_ (21.8 mg, 0.0669 mmol). After stirring at room temperature for 12 hrs (overnight), the reaction mixture was diluted with EtOAc (10 mL) and washed with water (5x10 mL), organic phase was dried (Na_2_SO_4_, and evaporated under vacuum. Crude product was purified by PTLC (DCM:MeOH:NH_4_OH, 90:9:1) to give 21 mg of product (58% yield). ^1^H NMR (400 MHz, DMSO-d6) δ 8.99 (s, 1H), 8.38 (t, 1H), 7.71 (d, J = 7.5 Hz, 1H), 7.68 – 7.55 (m, 2H), 7.55 – 7.43 (m, 1H), 7.34 (d, J = 7.8 Hz, 1H), 7.05 (s, 1H), 7.01 (d, J = 7.8 Hz, 1H), 5.11 (d, J = 3.9 Hz, 1H), 4.71 (d, J = 10.8 Hz, 1H), 4.51 (dd, 2H), 4.42 – 4.13 (m, 5H), 4.10 – 3.25 (m, 50H), 2.47 (s, 3H), 2.39 – 2.26 (m, 1H), 2.11 – 1.99 (m, 1H), 1.97 – 1.86 (m, 1H), 1.69 (p, J = 14.4, 6.9 Hz, 2H), 1.48 (p, J = 6.8 Hz, 2H), 1.43 – 1.22 (m, 4H), 0.97 (d, J = 6.4 Hz, 3H), 0.74 (d, J = 6.4 Hz, 3H). ^13^C NMR (151 MHz, dmso) δ 172.00, 168.53, 167.93, 156.30, 151.90, 148.35, 142.62, 132.02, 131.79, 131.69, 131.40, 128.34, 128.11, 127.60, 124.04, 123.45, 121.49, 112.55, 70.60, 70.51, 70.27, 70.24 (18C), 70.21, 69.92, 69.43, 69.05, 68.34, 59.14, 58.21, 55.84, 47.24, 45.80, 38.51, 37.50, 32.45, 29.48, 28.82, 26.54, 25.36, 19.30, 19.04, 16.44. HRMS (ESI); m/z: [M+H]^+^ Calcd. for C_59_H_92_ClN_4_O_17_S, 1195.5866. Found 1195.5869.

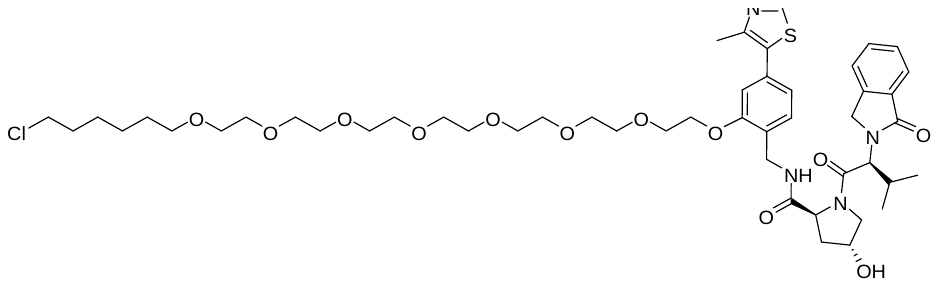

**(2S,4R)-N-(2-((27-chloro-3,6,9,12,15,18,21-heptaoxaheptacosyl)oxy)-4-(4-methylthiazol-5-yl)benzyl)-4-hydroxy-1-((S)-3-methyl-2-(1-oxoisoindolin-2-yl)butanoyl)pyrrolidine-2-carboxamide** (**SJF-7432**). To a mixture of(2S,4R)-4-hydroxy-N-[[2-hydroxy-4-(4-methylthiazol-5-yl)phenyl]methyl]-1-[(2S)-3-methyl-2-(1-oxoisoindolin-2-yl)butanoyl]pyrrolidine-2-carboxamide (25.3 mg, 0.0461 mmol) and 27-chloro-3,6,9,12,15,18,21-heptaoxaheptacosyl 4-methylbenzenesulfonate (**4**) (23.0 mg, 0.0384 mmol) in DMF (1 mL) was added Cs_2_CO_3_ (25.0 mg, 0.0768 mmol). After stirring at room temperature for 12 hrs (overnight), the reaction mixture was diluted with EtOAc (10 mL) and washed with water (5x10 mL), organic phase was dried (Na_2_SO_4_, and evaporated under vacuum. Crude product was purified by PTLC (DCM:MeOH:NH_4_OH, 90:9:1) to give 25 mg of product (67% yield). ^1^H NMR (500 MHz, DMSO-d6) δ 8.99 (s, 1H), 8.36 (t, J = 5.2 Hz, 1H), 7.71 (d, J = 7.4 Hz, 1H), 7.67 – 7.55 (m, 2H), 7.50 (t, 1H), 7.34 (d, J = 7.7 Hz, 1H), 7.05 (s, 1H), 7.01 (d, J = 7.8 Hz, 1H), 5.09 (d, J = 3.5 Hz, 1H), 4.72 (d, J = 10.7 Hz, 1H), 4.60 – 4.42 (m, 2H), 4.45 – 4.12 (m, 5H), 4.04 – 3.41 (m, 31H), 3.35 (t, J = 6.5 Hz, 2H), 2.47 (s, 3H), 2.34 (dq, J = 12.2, 6.6 Hz, 1H), 2.04 (dd, J = 12.7, 8.1 Hz, 1H), 1.92 (ddd, J = 12.9, 7.9, 4.7 Hz, 1H), 1.69 (dt, J = 13.1, 6.5 Hz, 2H), 1.47 (dt, J = 12.7, 6.4 Hz, 2H), 1.37 (dt, J = 14.2, 7.2 Hz, 2H), 1.34 – 1.22 (m, 2H), 0.97 (d, J = 6.1 Hz, 3H), 0.74 (d, J = 6.1 Hz, 3H).  ^13^C NMR (151 MHz, dmso) δ 171.53, 168.07, 167.45, 155.85, 151.45, 147.91, 142.19, 131.56, 131.37, 131.25, 130.96, 127.89, 127.67, 127.18, 123.61, 123.00, 121.05, 112.13, 70.16, 70.08, 69.84, 69.82, 69.80, 69.78 (6C), 69.48, 69.00, 68.61, 67.90, 58.70, 57.77, 55.41, 46.81, 45.36, 38.10, 37.06, 32.02, 29.05, 28.39, 26.11, 24.93, 18.88, 18.63, 16.03. HRMS (ESI); m/z: [M+H]^+^ Calcd. for C_49_H_72_ClN_4_O_12_S, 975.4555. Found 975.4532.

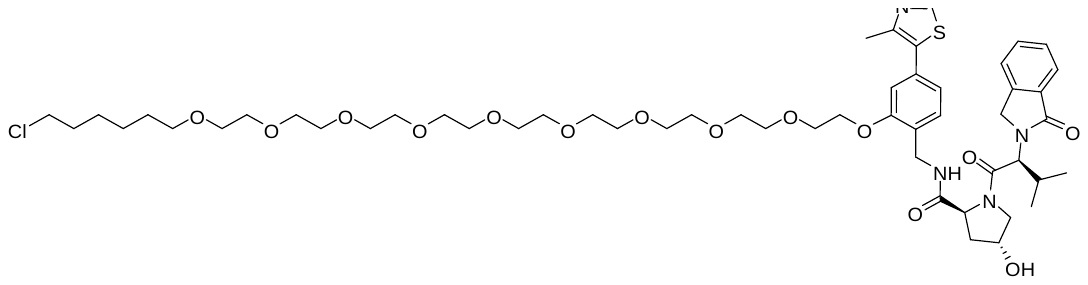

**(2S,4R)-N-(2-((33-chloro-3,6,9,12,15,18,21,24,27-nonaoxatritriacontyl)oxy)-4-(4-methylthiazol-5-yl)benzyl)-4-hydroxy-1-((S)-3-methyl-2-(1-oxoisoindolin-2-yl)butanoyl)pyrrolidine-2-carboxamide** (**SJF-7434**). To a mixture of (2S,4R)-4-hydroxy-N-[[2-hydroxy-4-(4-methylthiazol-5-yl)phenyl]methyl]-1-[(2S)-3-methyl-2-(1-oxoisoindolin-2-yl)butanoyl]pyrrolidine-2-carboxamide (22.0 mg, 0.0402 mmol) and 27-chloro-3,6,9,12,15,18,21-heptaoxaheptacosyl 4-methylbenzenesulfonate (**5**) (23.0 mg, 0.0335 mmol) in DMF (1 mL) was added Cs_2_CO_3_ (21.8 mg, 0.0669 mmol). After stirring at room temperature for 12 hrs (overnight), the reaction mixture was diluted with EtOAc (10 mL) and washed with water (5x10 mL), organic phase was dried (Na_2_SO_4_, and evaporated under vacuum. Crude product was purified by PTLC (DCM:MeOH:NH_4_OH, 90:9:1) to give 20 mg of product (56% yield). ^1^H NMR (500 MHz, DMSO-d6) δ 8.99 (s, 1H), 8.39 (t, J = 5.4 Hz, 1H), 7.71 (d, J = 7.5 Hz, 1H), 7.67 – 7.55 (m, 2H), 7.54 – 7.45 (m, 1H), 7.34 (d, J = 7.7 Hz, 1H), 7.05 (s, 1H), 7.01 (d, J = 7.8 Hz, 1H), 5.11 (d, J = 3.6 Hz, 1H), 4.71 (d, J = 10.8 Hz, 1H), 4.60 – 4.12 (m, 7H), 3.95 – 3.18 (m, 35H), 2.47 (s, 3H), 2.38 – 2.26 (m, 1H), 2.09 – 1.99 (m, 1H), 1.98 – 1.84 (m, 1H), 1.69 (p, J = 6.7 Hz, 2H), 1.47 (p, J = 6.6 Hz, 2H), 1.41 – 1.33 (m, 2H), 1.32 – 1.26 (m, 2H), 0.96 (d, J = 6.2 Hz, 3H), 0.73 (d, J = 6.4 Hz, 3H). ^13^C NMR (151 MHz, dmso) δ 171.54, 168.08, 167.46, 155.86, 151.46, 147.92, 142.19, 131.57, 131.37, 131.25, 130.97, 127.90, 127.68, 127.18, 123.61, 123.01, 121.06, 112.13, 70.17, 70.09, 69.85, 69.83, 69.81, 69.78 (17C), 69.74, 69.73, 69.49, 69.00, 68.62, 67.91, 58.71, 57.78, 55.42, 54.93, 46.81, 45.37, 38.10, 37.07, 32.03, 29.06, 28.40, 26.12, 24.94, 18.88, 18.63, 16.04. HRMS (ESI); m/z: [M+H]^+^ Calcd. for C_53_H_80_ClN_4_O_14_S, 1063.5080. Found 1063.5063

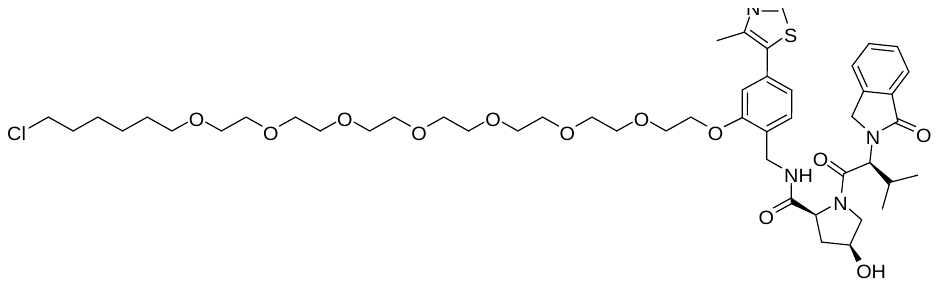

**(2S,4S)-N-(2-((27-chloro-3,6,9,12,15,18,21-heptaoxaheptacosyl)oxy)-4-(4-methylthiazol-5-yl)benzyl)-4-hydroxy-1-((S)-3-methyl-2-(1-oxoisoindolin-2-yl)butanoyl)pyrrolidine-2-carboxamide** (**JH-6073**). To a mixture of (2S,4S)-4-hydroxy-N-[[2-hydroxy-4-(4-methylthiazol-5-yl)phenyl]methyl]-1-[(2S)-3-methyl-2-(1-oxoisoindolin-2-yl)butanoyl]pyrrolidine-2-carboxamide (22.0 mg, 0.0402 mmol) and 2-[2-[2-[2-[2-[2-[2-[2-[2-(6-chlorohexoxy)ethoxy]ethoxy]ethoxy]ethoxy]ethoxy]ethoxy]ethoxy] ethoxy]ethyl 4-methylbenzenesulfonate (23.0 mg, 0.0335 mmol) in N,N-Dimethylformamide (1 mL) was added Cs_2_CO_3_ (21.8 mg, 0.0669 mmol). After stirring at room temperature for 12 hrs (overnight), the reaction mixture was diluted with EtOAcEtOAc (10 mL) and washed with water (5x10 mL), organic phase was dried (Na_2_SO_4_, and evaporated under vacuum. Crude product was purified by PTLC (DCM:MeOH, 9:1) to give 18 mg of product (56% yield). ^1^H NMR (500 MHz, cdcl3) δ 8.69 (s, 1H), 7.83 (d, J = 7.7 Hz, 1H), 7.62 (t, J = 6.1 Hz, 1H), 7.55 (t, J = 7.4 Hz, 1H), 7.46 (dt, J = 7.5, 3.6 Hz, 2H), 7.32 (d, J = 7.7 Hz, 1H), 6.99 (d, J = 7.7 Hz, 1H), 6.92 (s, 1H), 4.84 (d, J = 11.0 Hz, 1H), 4.76 (d, J = 17.5 Hz, 1H), 4.55 (d, J = 9.0 Hz, 1H), 4.50 (t, J = 5.6 Hz, 2H), 4.47 – 4.36 (m, 2H), 4.22 (q, J = 4.9 Hz, 2H), 4.09 (dd, J = 11.4, 4.4 Hz, 1H), 3.99 – 3.89 (m, 3H), 3.74 (q, J = 4.4 Hz, 2H), 3.68 – 3.54 (m, 26H), 3.52 (t, J = 6.7 Hz, 2H), 3.45 (t, J = 6.5 Hz, 3H), 2.53 (s, 3H), 2.41 – 2.03 (m, 4H), 1.77 (m, 5H), 1.59 (p, J = 6.9 Hz, 2H), 1.44 (t, J = 7.7 Hz, 2H), 1.45 – 1.31 (m, 2H), 0.89 (dd, J = 10.6, 6.5 Hz, 6H). ^13^C NMR (151 MHz, cdcl3) δ 172.68, 170.77, 169.14, 156.93, 150.52, 148.61, 142.17, 132.60, 131.93, 131.83, 129.88, 128.17, 126.35, 123.93, 123.03, 122.16, 114.64, 112.87, 71.37, 71.03, 70.94, 70.75, 70.72, 70.70, 70.67, 70.65, 70.24, 69.79, 68.05, 60.08, 58.33, 58.27, 47.26, 45.20, 39.55, 35.86, 32.68, 29.59, 29.05, 26.84, 25.56, 19.14, 18.91, 16.24. LC-MS (ESI); m/z [M+Na]^+^; Calcd. C_53_H_79_ClN_4_NaO_14_S, 1085.4900. Found 1085.4894.

**Materials and Methods**

**Cloning**

pCDNA5-CT-dCas9HT7 and pCDNA5-NT-dCas9HT7 constructs were generated by USER cloning strategy. For pCDNA5-CT-dCas9HT7, CT-dCas9HT7 ORF sequence was directly amplified using pET302-CT-dcas9HT7 plasmid as the template (Forward primer: 5’ACCTGACTATGCTGGAGTGGATAAGAAATACUCAATAGGCTTAGCTATCGGC3’ ; Reverse primer: 5’ATCAGCGGGTTTAACCGGAAATCUCCAGAGTAGACAGCC3’) and the pCDNA5 vector was amplified using pCDNA5 empty vector (Forward primer: 5’ AGATTTCCGGTTAAACCCGCTGAUCAGCCTCGAC3’; Reverse primer: 5’ AGTATTTCTTATCCACTCCAGCATAGTCAGGUACGTCATAAGGG3’). Amplified insert and vector PCR products were gen purified and subjected to USER cloning reaction using USER enzyme (NEB). Reaction mixture was then transformed into NEB5 cells and single colonies were sequenced to identify successful cloning products. To generate pCDNA5-NT-dCas9HT7 construct, a 2 step USER cloning strategy was employed. First, HaloTag7 sequence was amplified without the stop codon using pET302-CT-dcas9HT7 (Forward primer:5’ACGTACCTGACTATGCTGGAGCAGAAAUCGGTACTGGCTTTCCATTCG3’; Reverse primer: 5’ATCAGCGGGTACCGGAAATCUCCAGAGTAGACAGC3’) as the template and subjected to USER cloning reaction together with amplified pCDNA5 vector
(Forward primer:5’AGATTTCCGGTACCCGCTGAUCAGCCTCG3’; Reverse primer: 5’ ATTTCTGCTCCAGCATAGTCAGGTACGUCATAAGGG3’). In the second step, (Forward primer:5’ATTTCCGGTGGTGGCTCCAGAUCTGTGGATAAGAAATACTCAA

TAGGCTTAGCTATCGGC3’; Revisers primer: 5’ AGCGGGTTTAGTCACCTCCTAGCU

GACTCAAATCAATGC3’) dCas9 sequence was amplified with a stop codon and subjected to USER cloning reaction with the amplified vector obtained from the clone generated in the first step (Forward primer: 5’ AGCTAGGAGGTGACTAAACCCGCUGAT

CAGCCTCG3’; Reverse primer: 5’ ATCTGGAGCCACCACCGGAAAUCTCCAGAGTAG

ACAGC 3’). N terminal HA tag was introduced by the USER primers in both cases.

**Protein purification**

C-terminal HaloTag7-containing dCas9 fusion protein was expressed in *E. coli* BL21-RIPL codon plus bacterial cells. BL21 cells were transformed with 50 ng of plasmid DNA encoding dCas9HT7 and transformed cells were plated on carbenicillin-containing agar plates. On the following day, a single colony was selected and inoculated in 5 mL of LB and incubated overnight at 37 degrees. After 16 h, bacterial cells were diluted in 1 L of LB and shook at 37 degrees until OD600 reaches 0.8. Cells were kept on ice and induced with 0.5 mM IPTG and incubated 20 h at 18 degrees in a shaker. Cells were subjected to lysis (20 mM HEPES pH 8, 1mM MgCl2, 10% glycerol, 300 mM NaCl, 1 mM ß-ME and 1X protease inhibitor cocktail (Roche)) by exposing cells to 4 cycles of 30 seconds pulses and 1-minute rest on ice. Then clarified lysate was incubated with pre-washed Ni-NTA agarose beads (Agilent Technologies) for 1 h, at 4 degrees. The Ni-NTA beads were washed twice with wash buffer A and wash buffer B (Wash buffer A:20 mM HEPES pH 8, 1 mM MgCl2, 10% glycerol, 300 mM NaCl, 5 mM imidazole; Wash buffer B:20 mM HEPES pH 8, 1 mM MgCl2, 10% glycerol, 300 mM NaCl, 35 mM imidazole). Enriched dCas9HT7 protein was eluted with the elution buffer (20 mM HEPES pH 8, 1mM MgCl2, 10% glycerol, 300 mM NaCl, 300 mM imidazole). Eluted protein was further purified by gel filtration chromatography using a Superdex 200 column (GE Healthcare) in the storage buffer (20 mM HEPES pH 8, 1mM MgCl2, 300 mM NaCl). Purified protein was dialyzed against the storage buffer containing 10% glycerol. Purity of the dCas9HT7 protein was assessed by Coomassie staining.

**Annealing reaction**

Single stranded TRAFTACs and reverse oligonucleotides were dissolved in ultra-pure, RNAase free water. All the steps in this protocol were carried out in a clean, RNAase/DNAase-free environment. All the equipment and plasticware in this protocol were treated with RNAase Away prior to their use. Single stranded TRAFTACs and single stranded reverse oligonucleotides were mixed (final concentrations of TRAFTACs were set to 25 µM) in 1X annealing buffer (10 mM Tris, pH 7.5, 50 mM NaCl and 1 mM EDTA) and incubated for 5 minutes in a water bath at 95 degrees. Then, the hot-plate was turned off and the samples left to cool down to room temperature over 1-2 h. Double stranded TRAFTACs were mixed well, aliquoted and stored at -80 for the future use.

**EMSA**

Increasing concentrations of purified dCas9HT7 protein were incubated with or without 500 nM of NFκB-TRAFTAC or brachyury-TRAFTAC for 30 min at RT. Then the mixture was separated in a 1% agarose gel for 30 minutes at constant 120 mV and images were captured using ChemiDoc system (BioRad). For ternary complex formation assay, 3 µM of dCas9HT7 and increasing concentrations of purified brachyury was incubated with or without brachyury-TRAFTAC or control-TRAFTAC for 30 minutes at RT. Gel shifts were analyzed as described above.

**Cell culture**

Human embryonic kidney cells HEK293 cells and Hela cells were grown in Dulbecco’s Modified Eagles Medium (DMEM) containing 10% heat inactivated fetal bovine serum (FBS), 5 ug/mL streptomycin and 5 U/mL penicillin. All the cell culture experiments and maintenance were carried out in a humidified incubator at 37 degrees and 5% CO2 supplementation. One day prior to the transfection of chimeric oligos, cells (3.5X10^6^) were split into 6 cm cell culture dishes in complete growth medium. On the day of transfection cell culture medium was replaced with 3mL of transfection medium (DMEM 10%FBS in DMEM). Chimeric oligo transfection was performed using RNAi-Max according to the manufacturer’s protocol. Briefly, 25 nM of chimeric oligo and 15 µl of RNAi-Max were mixed in 125 µl of OPTI-MEM medium in two separate Eppendorf tubes. Mix two tubes together after 5 minutes and incubated for another 5 minutes prior to the addition to cells. After 6 h of transfection, the cells were split into 6-well plates and the transfection medium containing chimeric oligo was evenly divided into each well. Then cells were incubated for another 24 h and replaced with 1 mL of fresh transfection medium containing different concentrations of HaloPROTACs and incubated for 1 h prior to the addition of 0.25 mL of 5 ng/mL of TNF-alpha. Cells were then incubated for desired time at 37 degrees followed by cell lysis. Cell lysates were prepared by scraping off the cells using lysis buffer (25 mM Tris pH 7.4, 150 mM NaCl, 5 mM MgCl_2_, 1% NP40, 5% glycerol and 1X protease inhibitor cocktail from Roche) and lysates were centrifuged at high speed (14 000 rpm) for 10 minutes and clear supernatant was collected for further experiments.

**Immunoprecipitation**

Cells that overexpress dCas9HT7 together with either p50 or brachyury-GFP was lysed lysis buffer (25 mM Tris pH 7.4, 150 mM NaCl, 5 mM MgCl_2_, 1% NP40, 5% glycerol and 1X protease inhibitor cocktail from Roche) or IP buffer (25 mM Tris pH 7.4, 150 mM NaCl, 0.4% NP40, 5% glycerol and 1X protease inhibitor cocktail from Roche) respectively. Lysates (1 mg for each sample) were incubated with 100 nM of NFκB-TRAFTAC, brachyury-TRAFTAC or 3’controlcrRNA for 1 h, at RT. Then cell lysates mixture was incubated with 25 ul of pre-washed HA-agarose beads (Sigma) and top up to 500 ul with 1X TBS. Then tubes were incubated at 4 degrees overnight in a rotator. Tubes were centrifuged at 2000 rpm for 2 minutes at 4 degrees and beads were washed three times with 1 mL of lysis buffer or IP buffer for 3 times with 5 minutes incubation in a rotisserie. Beads were then eluted with 2X loading buffer containing 10% ß-ME and eluted samples were analyzed by western blotting by probing with desired primary antibodies. Chemiluminescence signal was captured using ChemiDoc system by BioRad.

**Immunofluorescence**

Cells were split into 8-well imaging slides one day prior to the transfection. On the day of transfection, cell culture medium was replaced with transfection medium and transfection of fluorescein-labelled chimeric oligo was carried out as described above. After 12 h of transfection, cells were fixed with 4% paraformaldehyde for 10 minutes followed by three washes with 1X PBS. Then cells were permeabilized with 0.1% Triton-X-100 in PBS for another 10 minutes. Cells were blocked for 1 h, at RT prior to the overnight incubation of anti-HA antibody. Cells then washed three times with PBS and secondary antibody conjugated to Alexa Fluor-567 was incubated for 1h at RT. After 3 washings with PBS cells were analyzed by confocal microscopy (Zeiss LSM 880 Microscope). For brachyury-GFP, images were directly captured using fluorescence microscope (EVOS M5000 IMAGING SYSTEM) without prior fixation or permeabilization of the cells.

**Supporting Figures**

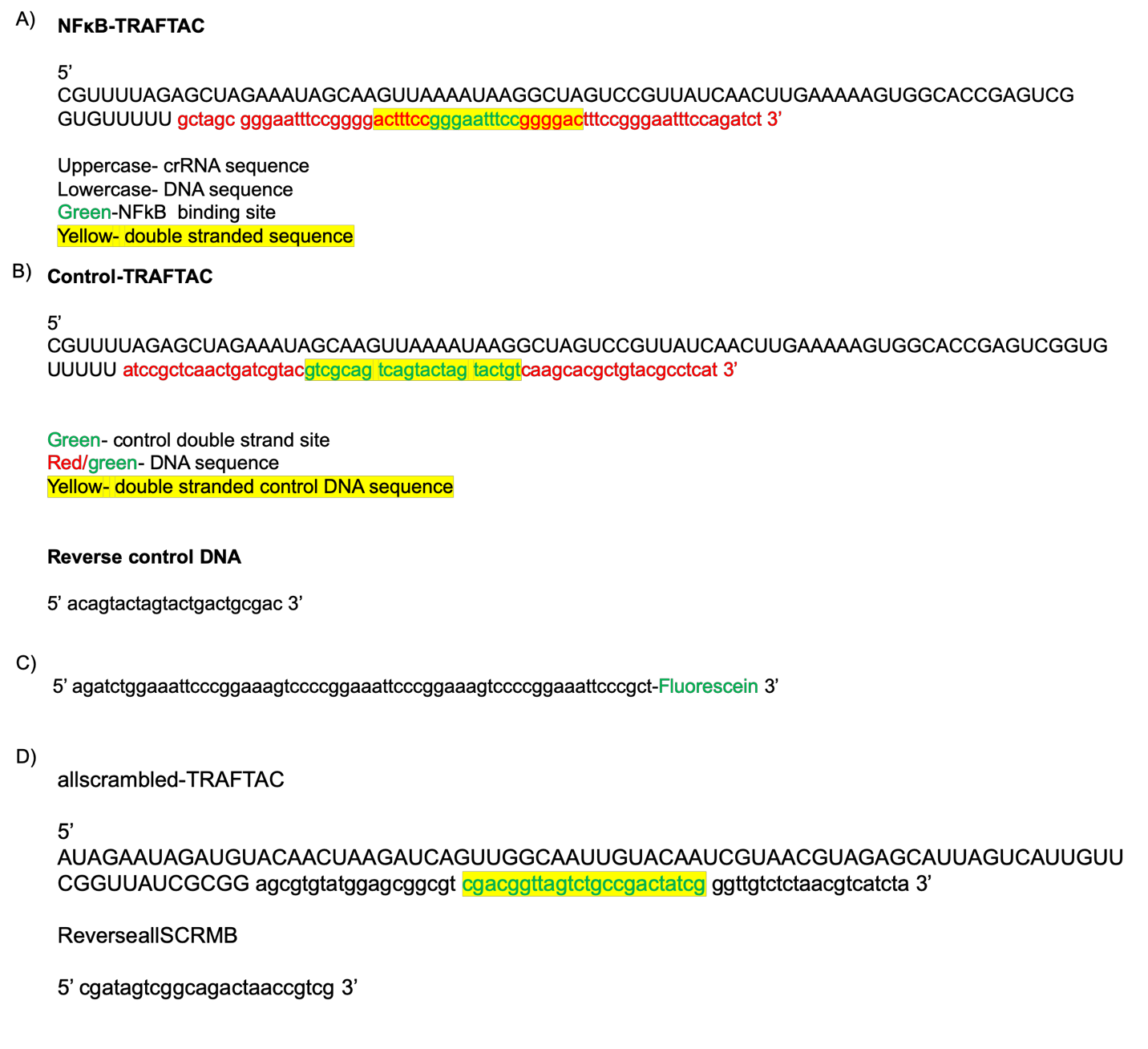

**Figure S1. Oligonucleotide sequences for TRAFTACs and reverse complements used in this study.** A) single stranded NFκB-TRAFTAC sequence. B) Single stranded control-TRAFTAC sequence. C) Reverse complement sequence covalently attached to fluorescein at the 5’ end of the sequence. D) Single stranded sequence of allscrambled-TRAFTAC.

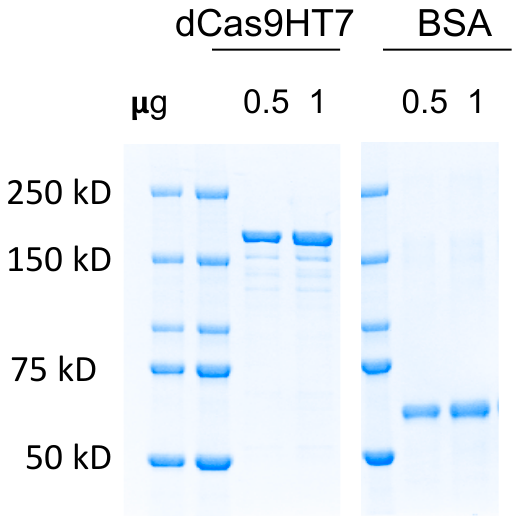

**Figure S2.** A fusion of HT7 and dCas9 proteins was purified using HIS-tag affinity purification and further purified by size exclusion chromatography. Purified fusion dCas9HT7 protein was ran in parallel with BSA.

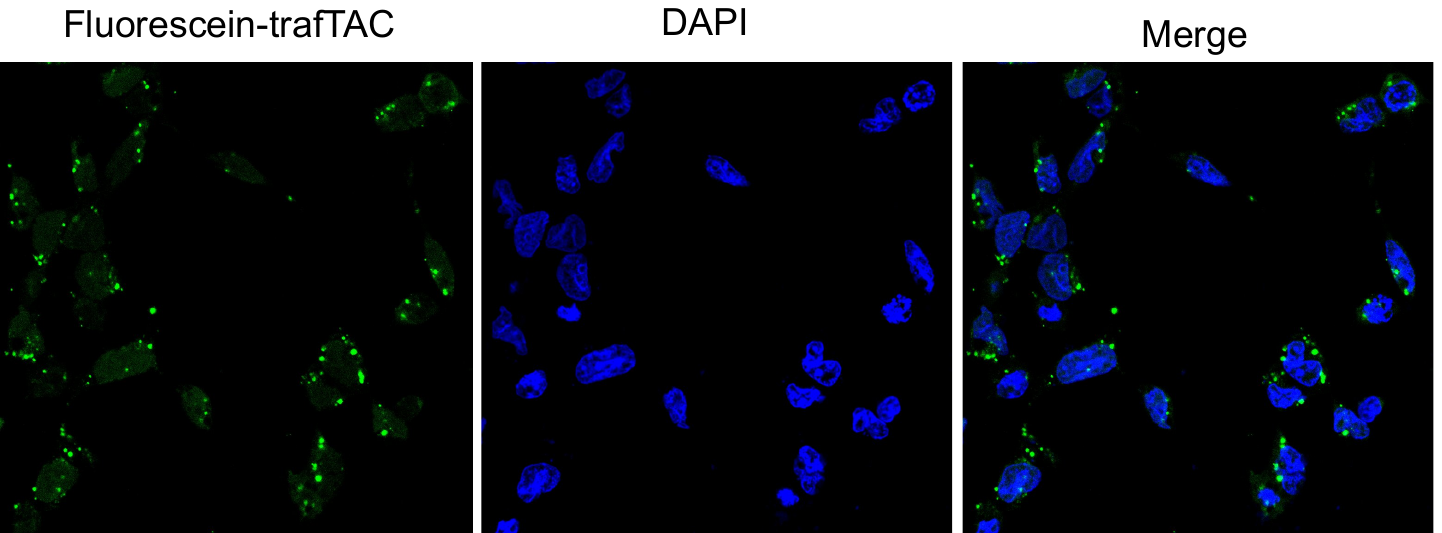

**Figure S3.** HEK293 cells were transfected with 25 nM of fluorescein-labeled TRAFTAC and after 12 h cells were fixed, permeabilized and labeled with DAPI. Cells were analyzed for fluorescein and DAPI signal by confocal microscopy at 40X magnification.

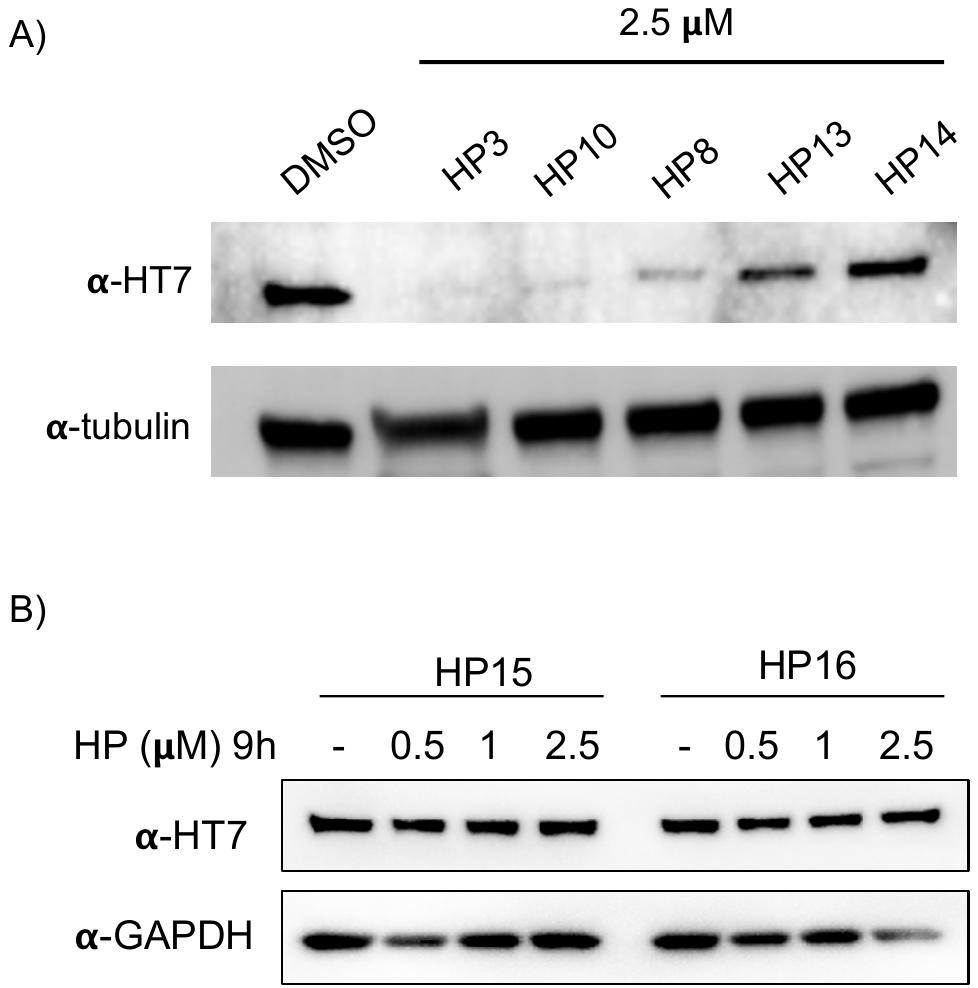

**Figure S4.** A) and B) HaloPROTAC screening towards the degradation of HaloTag fusion protein, dCas9HT7.

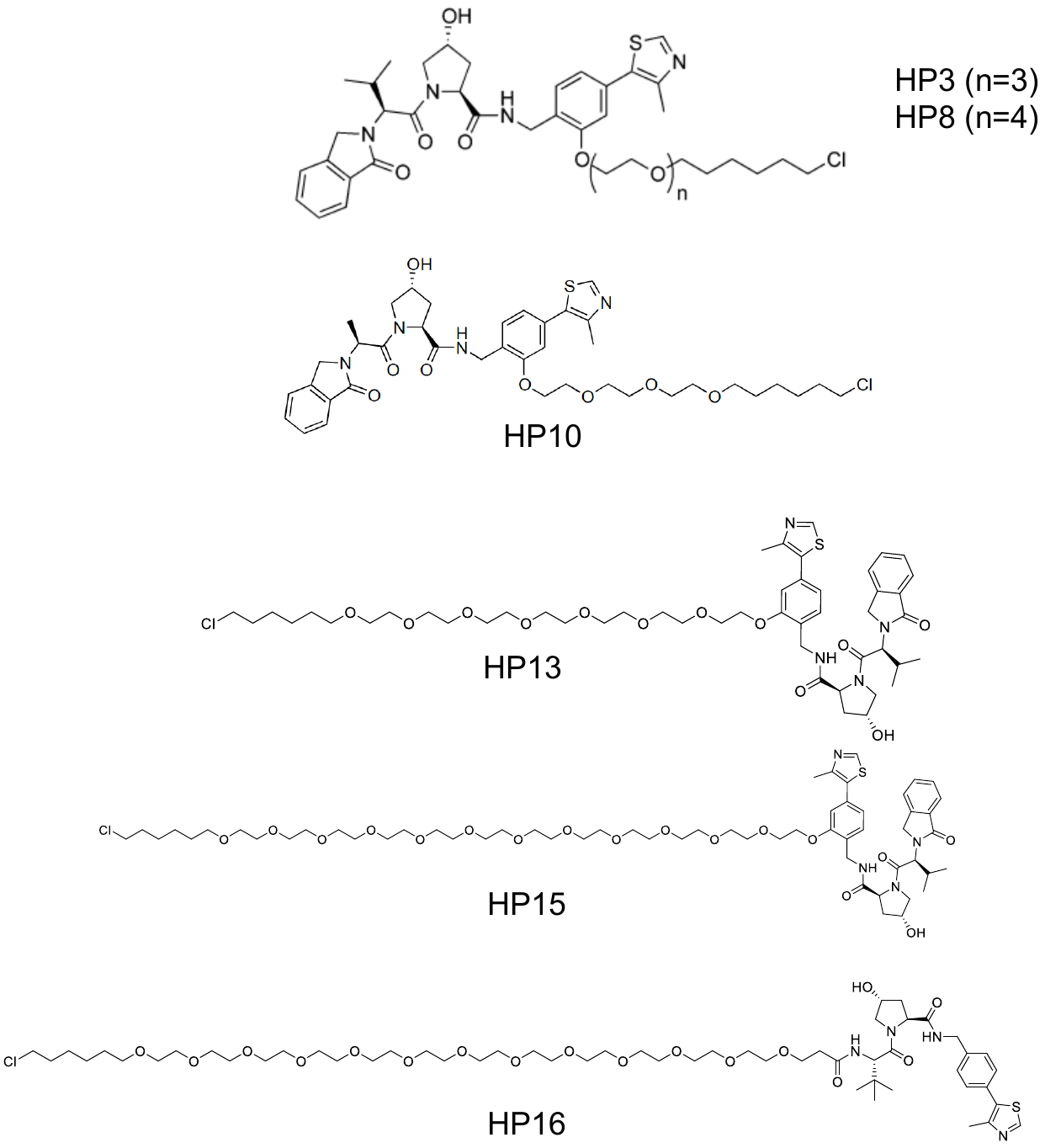

**Figure S5**. Chemical structures of HaloPROTACs (HP) used in this study.

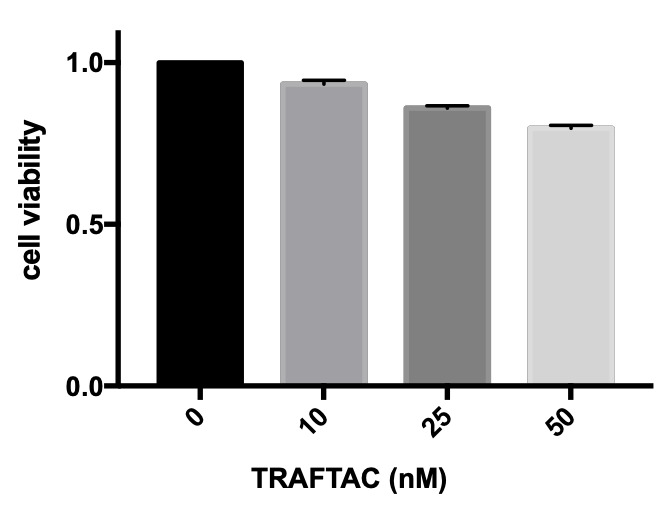

**Figure S6. Cell viability assay**. Stable cells that express dCas9HT7 were transfected with increasing concentrations of NFκB-TRAFTAC and after 36 h, cells were subjected to MTS assay.

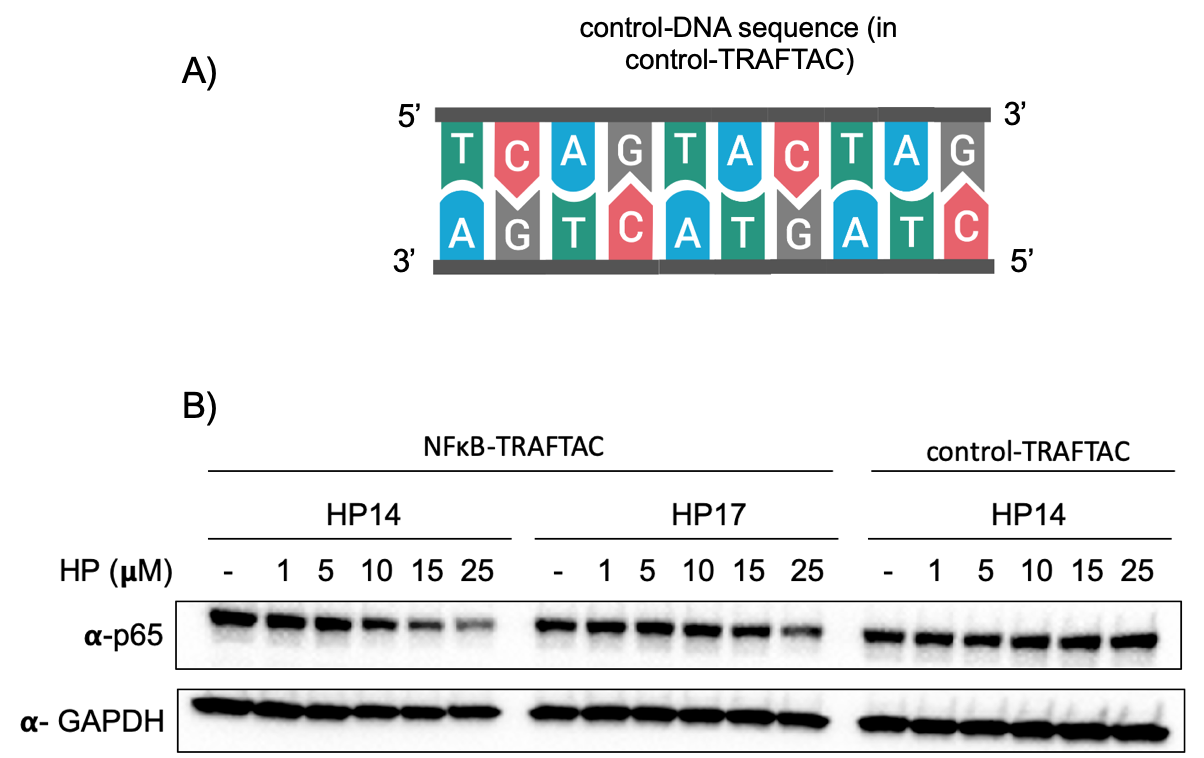

**Figure S7.** A) Double stranded, control oligonucleotide sequence used in the control-TRAFTAC. B) Stable cells overexpressing CT-dCas9HT7 were transfected with NFκB-TRAFTAC or control-TRAFTAC followed by HP and TNF-alpha treatment. Cells were lysed and analyzed for p65 and GAPDH levels.

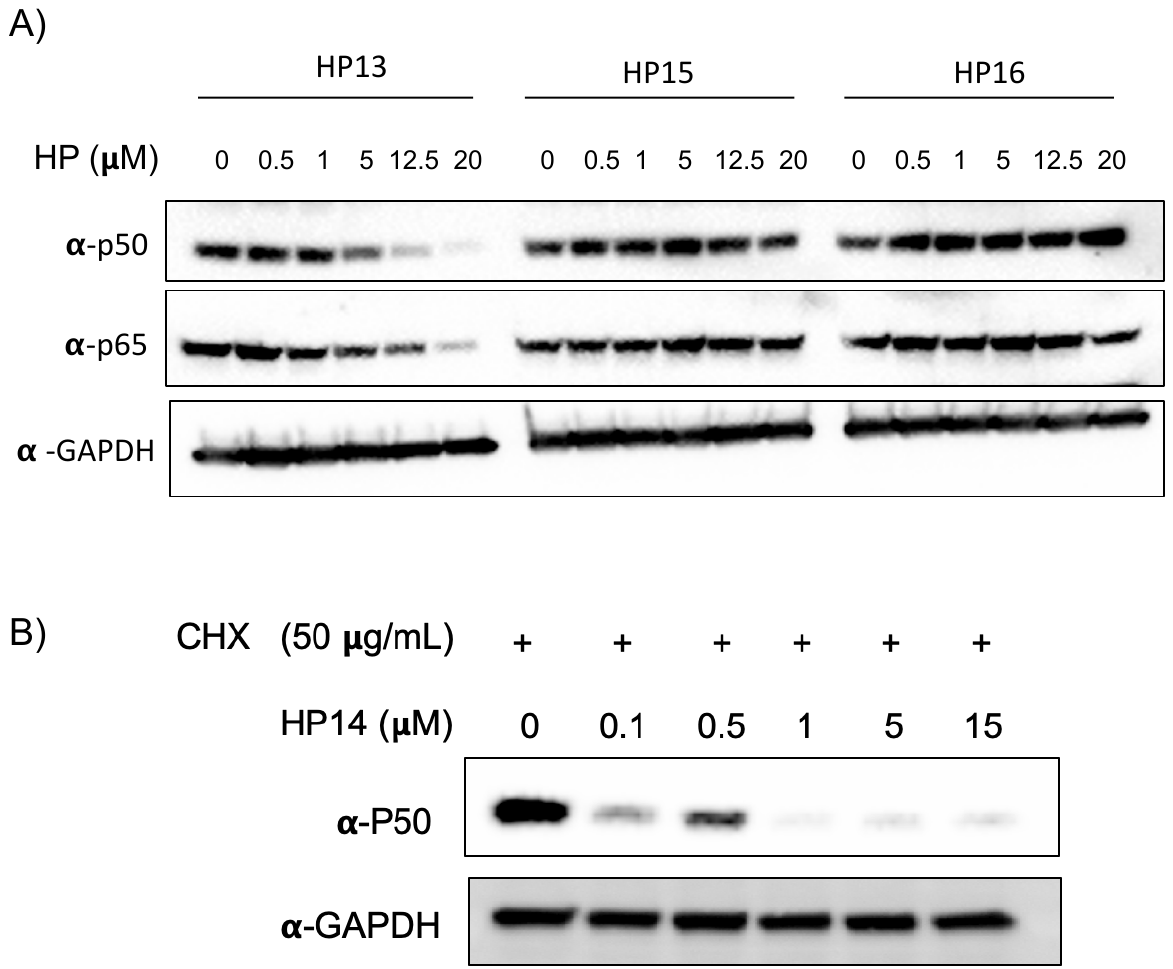

**Figure S8.** A) NFκB degradation by HP13, HP15 and HP16**.** Stable cells were transfected with NFκB-TRAFTAC followed by HP and TNF-alpha treatment. Cell lysates were probed as indicated. B) Cycloheximide co-incubation with HP14/ TNF-alpha, in NFκB-TRAFTAC transfected cells, significantly induced p50 degradation levels.

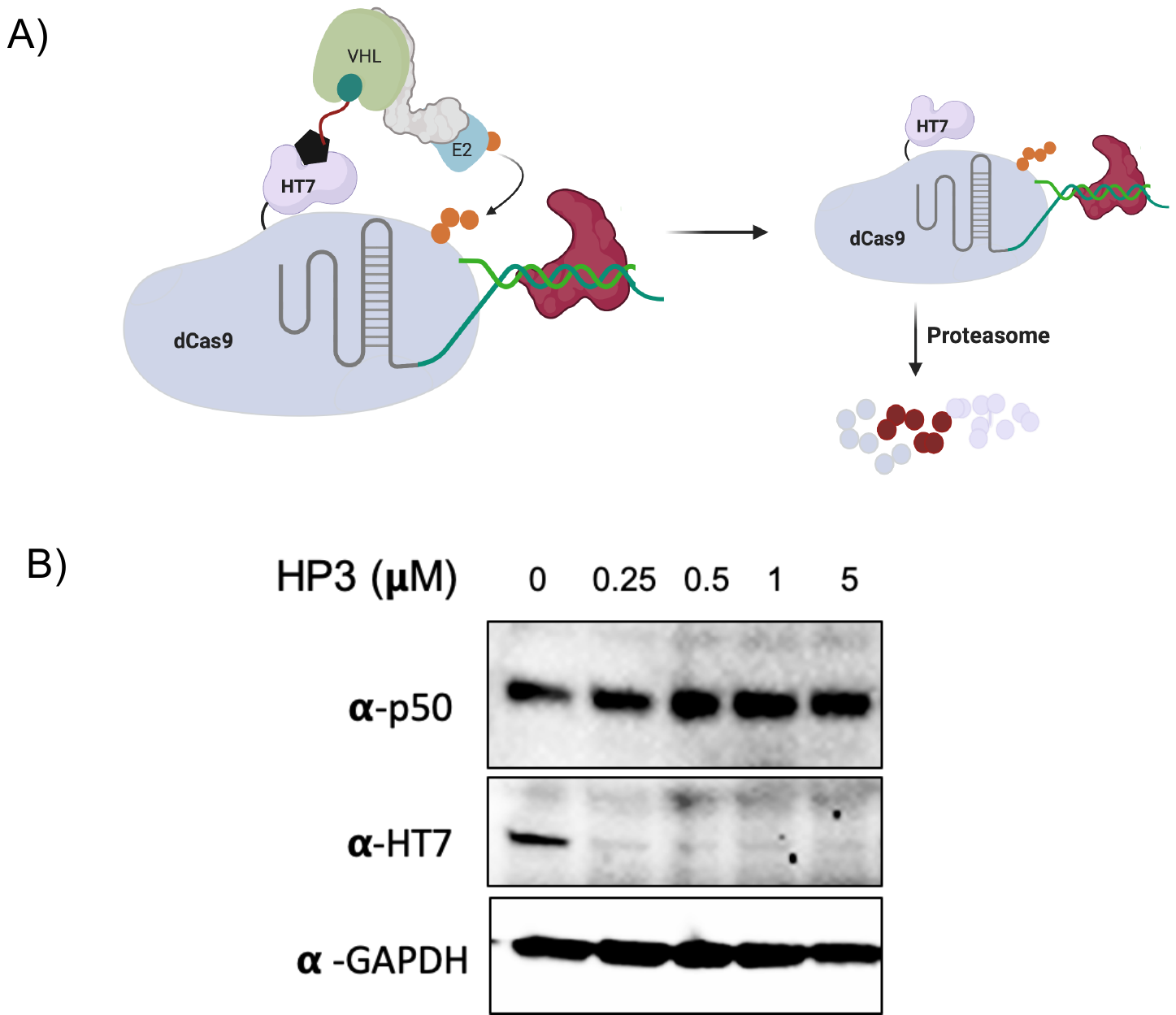

**Figure S9.** A) Schematic representation of possible HP3 mediated degradation of dCas9HT7:TRAFTAC:p50 complex. B) Stable cells were transfected with NFκB-TRAFTAC and treated with HP3 followed by TNF-alpha. Lysates were probed as indicated in the figure.

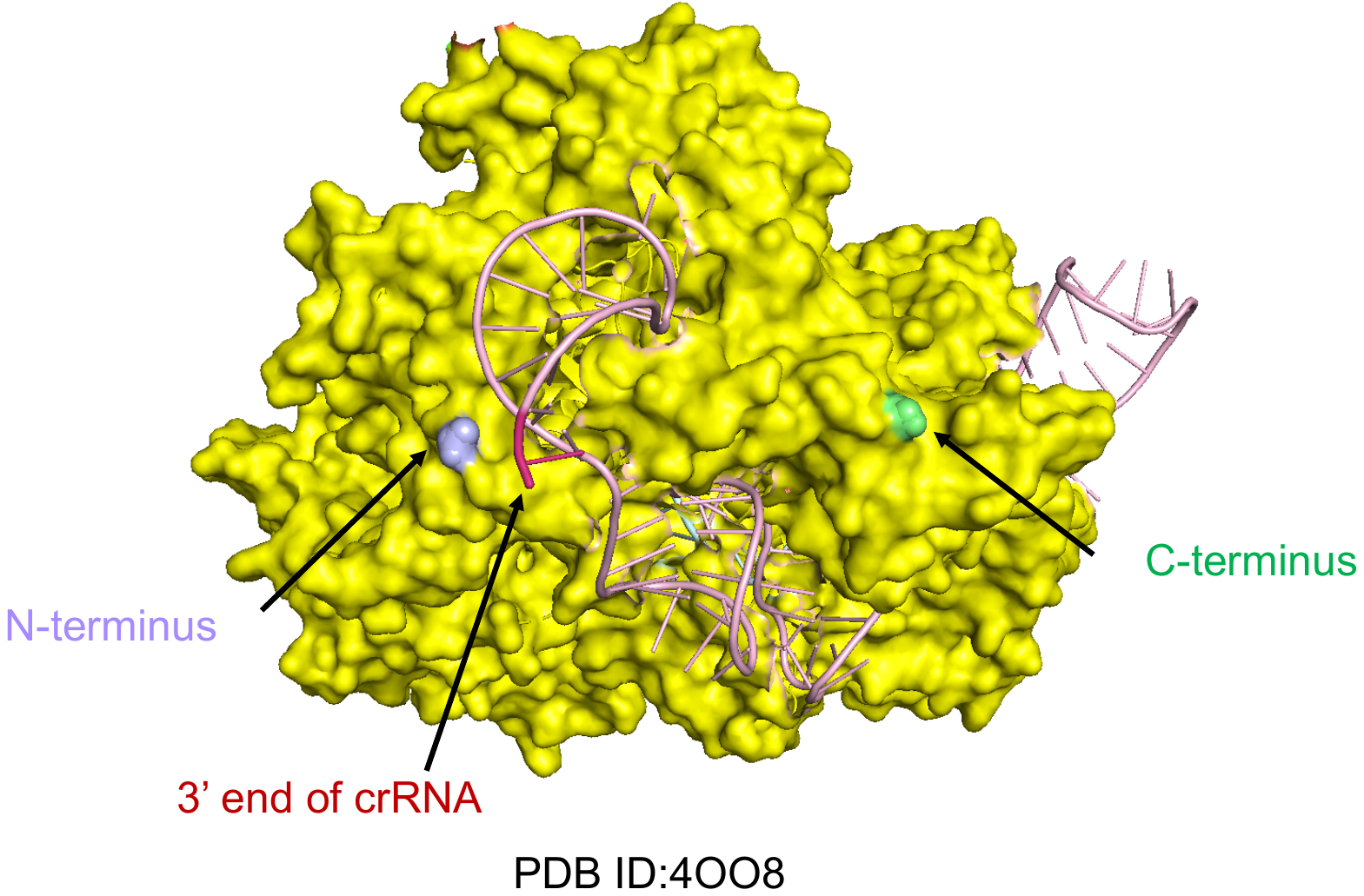

**Figure S10.** Crystal structure of *S. pyogenes* Cas9 in complexed with gRNA (PDB ID:4OO8). Both N- and C-terminal amino acids of the Cas9 protein are located in the space such that N- or C-terminally tagged fusion HT7 can access the 3’ end of the Cas9 bound gRNA.

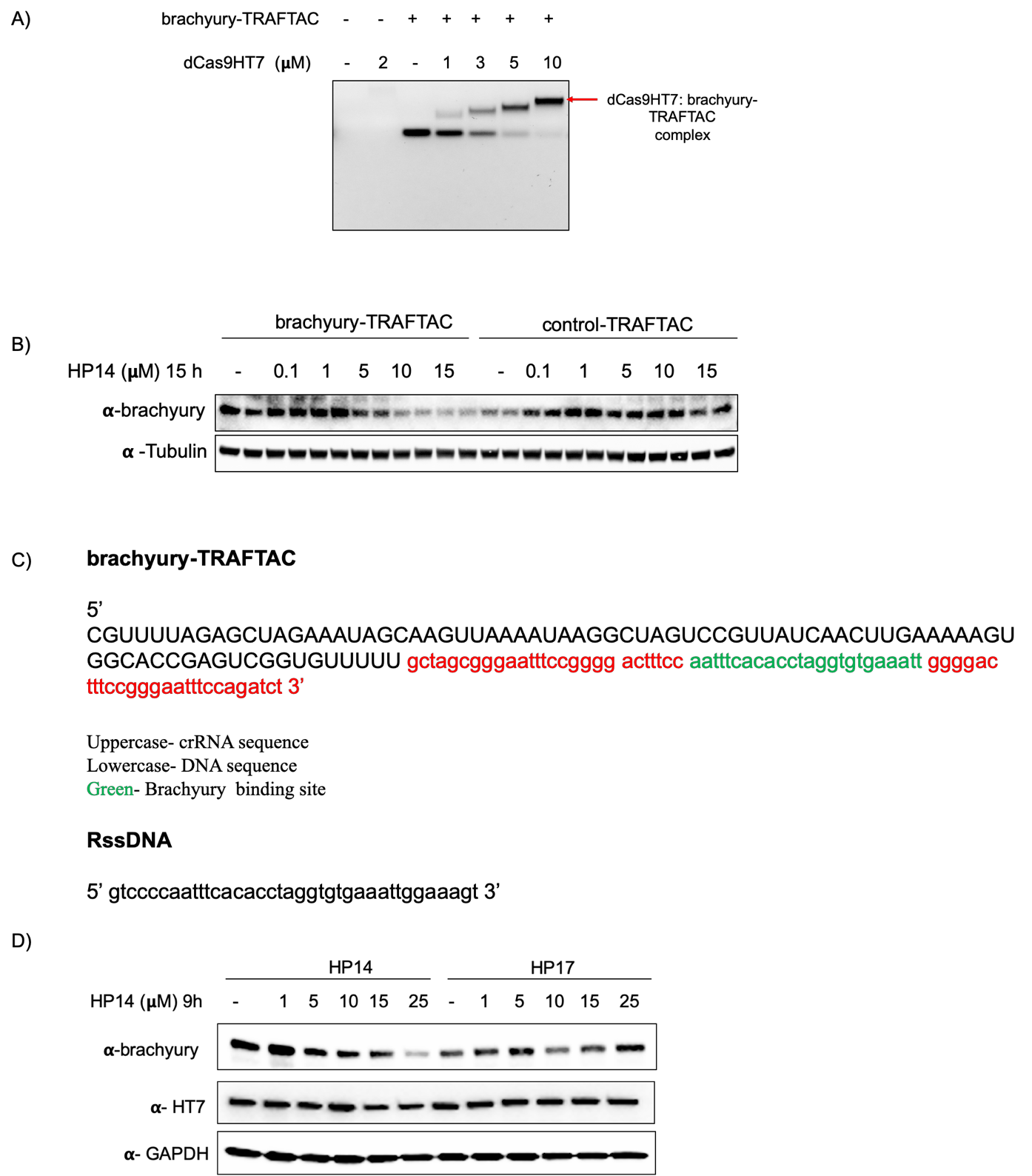

**Figure S11.** A) Fusion protein of dCas9HT7 forms the binary complex with brachyury-TRAFTACs *in vitro* as seen by EMSA. B) Cells were transfected with brachyury-TRAFTAC or control-TRAFTAC. Then cells were treated with HP14 for 15 and cell lysates were subjected to western blot analysis as shown in the figure. C) The chimeric oligo sequence of the single stranded brachyury-TRAFTAC and reverse complement sequence of brachyury targeting DNA sequence (shown in green). D) After transfection of brachyury-TRAFTAC, cells were treated with HP14 and HP17 for 9 h before cell lysis. Cell lysates were analyzed by western blot using antibody against brachyury, HT7and GAPDH.

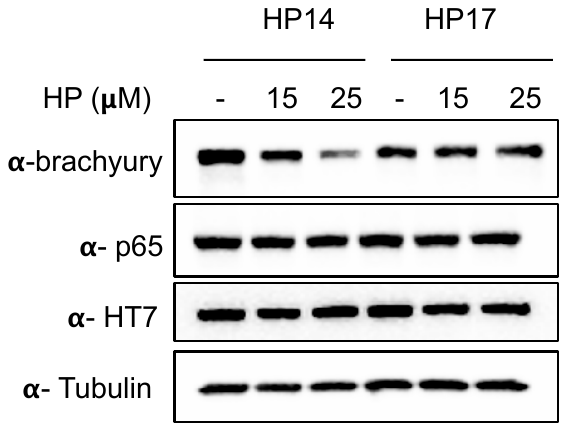

**Figure S12.** Brachyury targeting TRAFTAC did not induce degradation of other transcription factors. Cells were transfected with brachyury-TRAFTAC followed by HP14 and HP17 treatment. After 9 h, cells were lysed and lysates were analyzed for brachyury-GFP, p65, HT7 and tubulin.

**References**

1.	Maniaci C, Hughes SJ, Testa A, et al. Homo-PROTACs: bivalent small-molecule dimerizers of the VHL E3 ubiquitin ligase to induce self-degradation. *Nat Commun.* 2017;8(1): 830.
